# *Leptospira interrogans* triosephosphate isomerase: exploring the structural determinants of stability, high reaction rate and specificity

**DOI:** 10.1101/2020.12.04.412312

**Authors:** Vidhi Pareek, Vaigundan Dhayabarn, Hemalatha Balaram, Patnam R Krishnaswamy

## Abstract

Leptospires are zoonotic pathogens that cause significant socio-economic burden in developing countries, world-wide. The pathogenic species *Leptospira interrogans* (*Li*) is an important and interesting target for investigating the enzymes essential to its metabolic needs and adaptations. We cloned and expressed triosephosphate isomerase (*Li*TIM), a central metabolic flux regulator of *Li*, in AA200, *E. coli* TIM null strain. *Li*TIM was obtained as an active dimer (D-GAP→DHAP *k*_cat_ = 1740 s^−1^ and *K*_m (D-GAP)_ = 0.21 mM, at 25 °C) with mid-transition concentrations, C_m_, 0.8 mM and 2.6 mM, respectively, for guanidine hydrochloride and urea induced equilibrium unfolding. We report the high resolution X-ray structures of *Li*TIM in *apo* and substrate (DHAP) bound forms. Our analysis highlights key features of TIM that regulate the mode of substrate binding and transition state stabilization and thus play a decisive role in attainment of high proficiency for the isomerisation reaction while avoiding the elimination reaction. Unexpected differences in the effect of temperature on stability and activity were observed for the three mesophilic pathogenic TIMs *viz*. from *Li, Plasmodium falciparum* (*Pf*) and *Trypanosoma brucei* (*Tb*). *Li*TIM and *Tb*TIM (T_m_ = 46.5 °C) were more susceptible to unfolding and precipitation compared to *Pf*TIM (T_m_ = 67 °C). In contrast, the initial (or zero point) activity of *Pf*TIM rises till 50 °C and saturates unlike *Li*TIM and *Tb*TIM which show a rise till 55 °C and 60 °C, respectively. These observations could be rationalized by sequence comparison and examination of the structures of the three TIMs.

## INTRODUCTION

*Leptospira* are important zoonotic pathogens with ability to infect not just humans but livestock and other vertebrates of economic value [1–3], more so in large parts of the developing and underdeveloped countries [2, 4–6]. *Leptospira* are gram negative, obligate aerobic bacteria of the genus spirochaete [2, 7] and according to a recent estimate, the pathogen causes 1.03 million infections and 58, 900 human deaths world-wide per annum and 2.90 million disability adjusted life years (DALY) [8, 9]. In addition, it also infects cattle, rodents, horses, marsupials, dogs, sheeps and pigs [2, 6, 8]. While classified as a zoonotic pathogen of global importance by the WHO [3, 9, 10], leptospirosis is a comparatively neglected infection and the current understanding of the organism’s metabolism and the difference between pathogenic and non-pathogenic strains is fairly limited [10, 11]. How the metabolism of *Leptospira interrogans*, one of the pathogenic species, assists in adapting to different host environments is an intriguing biochemical puzzle and the solution might hold the key to developing control measures against its spread. Interestingly, *Leptospira* does not use glucose as its major carbon and energy source [12] though, genome annotations [13–15], proteomic analysis [16, 17] and *in vitro* recombinant enzyme characterization [18] have identified functional glycolytic pathway enzymes [19]. The bacterium has a functional TCA cycle and electron transport chain [20] and uses beta-oxidation of long chain fatty acids as the primary carbon source [21]. Current limitations in growing the parasite under laboratory conditions make diagnosis and studies difficult [5, 22]. Approaches to inhibitor design would benefit from the *in vitro* biochemical and structural analysis of the essential metabolic enzymes. From the published genome [13], we identified one such central metabolic enzyme of this pathogen, triosephosphate isomerase (TIM), which connects the fatty acid and glycerol metabolism to gluconeogenesis and TCA cycle via an anaplerotic route and provides intermediates essential for energy production and amino acid biosynthesis.

Triosephosphate isomerase (TIM; EC: 5.3.1.1) interconverts dihydroxyacetone phosphate (DHAP) and D-glyceraldehyde-3-phosphate (D-GAP), the triosephosphates that result from the breakdown of six-carbon sugars in glycolysis. TIM is a remarkable molecular catalyst, arguably a ‘perfect enzyme’ [23, 24], and achieves 9 orders of magnitude increment in the rate of isomerization of DHAP and GAP, compared to the uncatalyzed and acetate catalysed rates [25–27]. The TIM catalysed reaction proceeds via formation of an enediol intermediate. The intermediate may potentially interconvert DHAP and GAP by isomerisation or produce methylglyoxal (MG) and inorganic phosphate (Pi) by elimination [25, 28] (as in the enzyme methylglyoxal synthase [29, 30]), depending on the specific proton transfer steps [31]. It is remarkable that TIM catalyses isomerisation with high efficiency as well as specificity, avoiding the elimination reaction [32]; elimination from the enediol intermediate was found to be 100 times faster than isomerization during the uncatalyzed reaction in solution [25].

To investigate the functional relevance of *Li*TIM and its suitability as a drug target, we begun by cloning and characterization of the enzyme. We determined the biophysical and biochemical properties of *Li*TIM and compared that with two well-studied pathogenic TIMs *viz*., from *Trypanosoma brucei* (*Tb*TIM) and *Plasmodium falciparum* (*Pf*TIM). We determined the high resolution X-ray structure of *Li*TIM in its *apo* and *holo* (with the substrate DHAP) forms. The similarities and differences in the properties of *Li*TIM, *Tb*TIM and *Pf*TIM suggest that sequence variations at non-catalytic peripheral residues have differential effects on structural stability and dynamics essential for activity. The *Li*TIM substrate bound structure was of particular interest for gaining insights into the mechanistic features of TIM which continue to attract attention.

## RESULTS

### Cloning, expression and purification of *Li*TIM

Triosephosphate isomerase gene was amplified from *Leptospira interrogans* cDNA and cloned in the expression vector pTrc99A (primers given in Table S1). The protein was expressed in AA200, *tpi* null *E. coli* expression strain (yield ~ 25 mg/L, Figure 1A), and after the final purification step 95% pure protein was obtained as a well folded (circular dichroism spectrum (Figure 1B) and intrinsic tryptophan fluorescence, Figure 1C) dimer in solution (mono dispersed peak in analytical gel filtration, Figure 1E). The far-UV CD spectrum of *Li*TIM (from 200-250 nm) was similar to the reported spectra for TIMs from other organisms and characteristic of the canonical (β/α)_8_ fold [48, 49]. The spectrum shows one prominent negative band at ~208 nm and another very broad but more intense negative band at ~222 nm (likely including the contribution due to β-strands as well as helices). Additionally, aromatic side chains of Tyr, Phe and Trp may contribute to CD intensities around 222 nm [50, 51]. *Li*TIM has two Trp (positions 11 and 172), four Tyr and thirteen Phe residues (Figure 1F). The purified protein showed broad fluorescence emission band with λ_max_em = 325 nm when excited at 295 nm wavelength light. *Post facto* examination of the *Li*TIM structure revealed that while W11 lies in a hydrophobic environment and the ring −NH makes a hydrogen bond to one of the water molecules in the crystal structure, W172 is a part of loop-6 and is a solvent exposed tryptophan. The LC/ESI-MS of the purified protein confirmed its exact mass with loss of the N-terminal methionine (observed mass: 27177.9 Da, Figure 1G). The purified protein was used for kinetic and biophysical characterization and crystallization (Figure 1D and Table S2).

**Figure 1.**
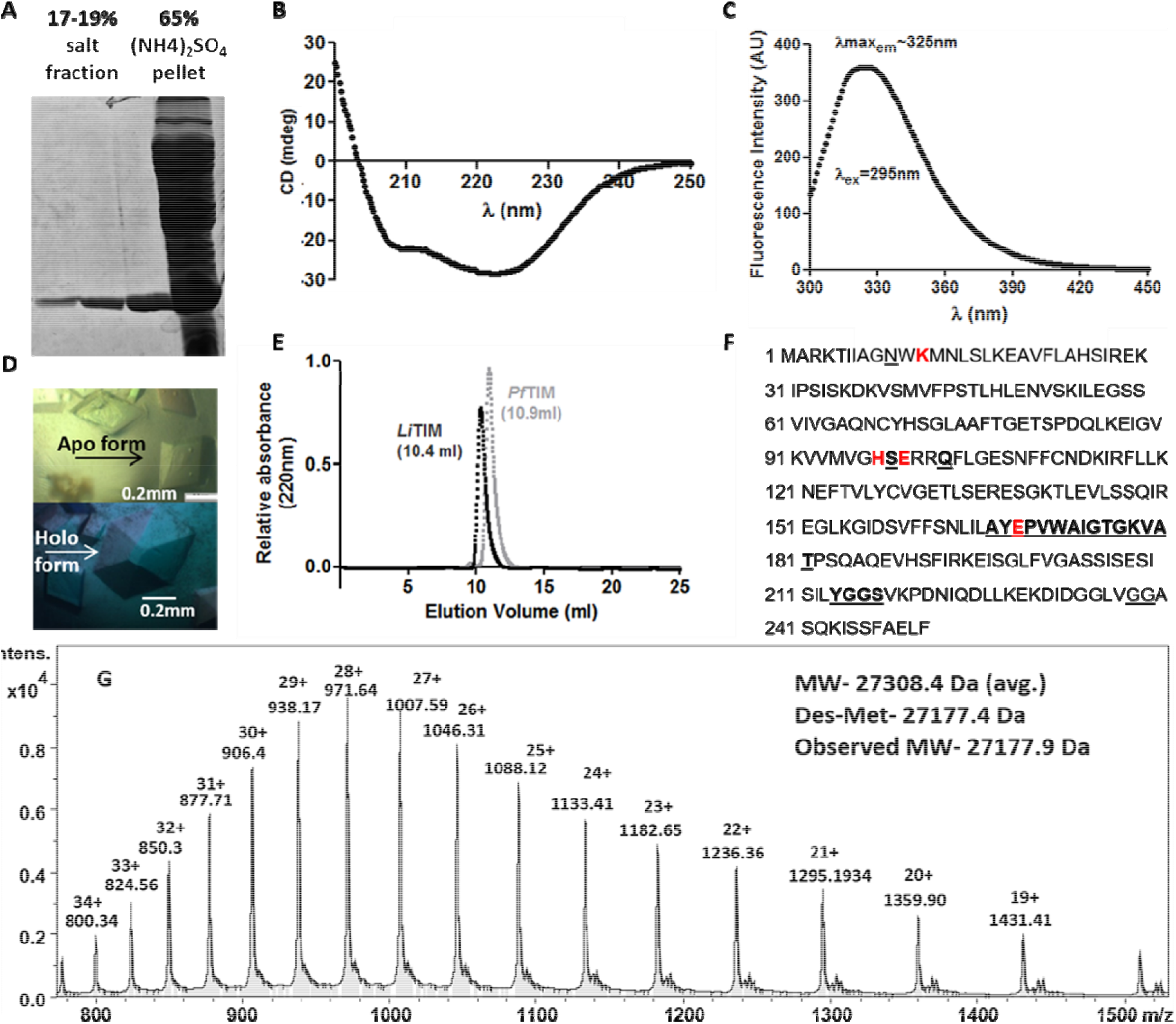
Purification and characterization of Leptospira interrogans TIM (LiTIM). (**A**) 12% SDS-PAGE showing the final purified protein used for further characterisation. (**B**) Far-UV CD spectrum and (**C**) tryptophan intrinsic fluorescence spectrum of the purified protein confirm a well folded form in solution. (**D**) Crystals obtained for the apo form of LiTIM. The crystal in the lower panel was soaked with substrate to obtain the holo form of the protein. (**E**) Analytical gel filtration chramatography indicates that it is a dimer in the solution (PfTIM WT, known to be dimeric, was used as the control [56]). (**F**) LiTIM protein sequence (251 residues)-catalytic residues in red bold, residues that showed conformational change upon substrate binding in bold underline, and other periferal, conserved residues that interact with substrate are shown in underlined letters. (**G**) LC-ESI/MS of the purified LiTIM protein agrees with the expected mass.

### Kinetic characterization

To ascertain the functional identity of the expressed recombinant protein, we determined the rate of isomerisation of D-GAP to DHAP using a continuous, coupled enzyme assay (details in Experimental Procedures section). Initial assays showed that upon prior incubation of diluted *Li*TIM solutions (~66 ng/ml) on ice, the protein lost activity (Figure 2A). Also, a non-linear increase of specific activity with rise in temperature from 35 to 45 °C was noticed (Figure 2B). Such loss of activity upon dilution of protein and incubation could either be because of dimer dissociation and precipitation or due to sticking of the protein on the walls of the container due to exposed hydrophobic surfaces. The ΔG dissociation for *Li*TIM dimer is ~−20 kcal/mol, as estimated using the dimer interface interactions and the total buried surface area (1750 Å^2^/monomer, PISA program, CCP4i) giving a dimer dissociation constant of ~2.8×10^−15^ M at 25 °C. Since we used a concentration of around ~ 6×10^−9^ M in our dilution mix and ~2.4×10^−10^ M in the final assay mix, dimer dissociation seems unlikely. To check the second possibility, we incubated the *Li*TIM protein with 1.5 μM BSA in the dilution mix as well as the reaction mix (BSA alone showed no background activity, data not shown). With the same concentration of *Li*TIM as the earlier assays, in the presence of BSA, the specific activity remained constant upto 60 minutes and increased linearly till 45 °C (Figure 2A and B). Using this method, D-GAP titration was performed at 25 °C and the specific activity values were fit to the Michaelis-Menten equation to obtain *k*_cat_ = 1700 s^−1^ and the *K*_m_ = 0.2 mM (Figure 2C; Table 1). We also determined the kinetic constants for two of the extensively studied pathogenic mesophilic TIMs *viz*., *Trypanosoma brucei* TIM, *Tb*TIM, (*k*_cat_ = 945 s^−1^; *K*_m_ = 0.17 mM) and *Plasmodium falciparum* TIM, *Pf*TIM, (*k*_cat_ = 500 s^−1^; *K*_m_ = 0.9 mM) under the same assay conditions at 25 °C (Table 1). The *Pf*TIM and *Tb*TIM kinetic parameters were in agreement with previous studies [33, 52]. Pairwise sequence alignment (ClustalOmega online server [53]) of the three protein sequences (SwissProt online database [54]) showed that the sequence identity between *Li*TIM-*Tb*TIM is 35%, *Li*TIM-*Pf*TIM is 40% and *Pf*TIM-*Tb*TIM is 42% out of which around 28% residues are identical between all the three, (Figure S2). While all the three enzymes are categorised as *mesophilic*, belong to the same fold class, have the same oligomeric state and were found to be active (Table 1) and stable at 25 °C, they are markedly different in residue composition. To understand the effect of peripheral residues on functional features of TIM, we investigated the effect of chaotropes and temperature on the stability and the effect of temperature on the activity of the three enzymes.

**Figure 2.**
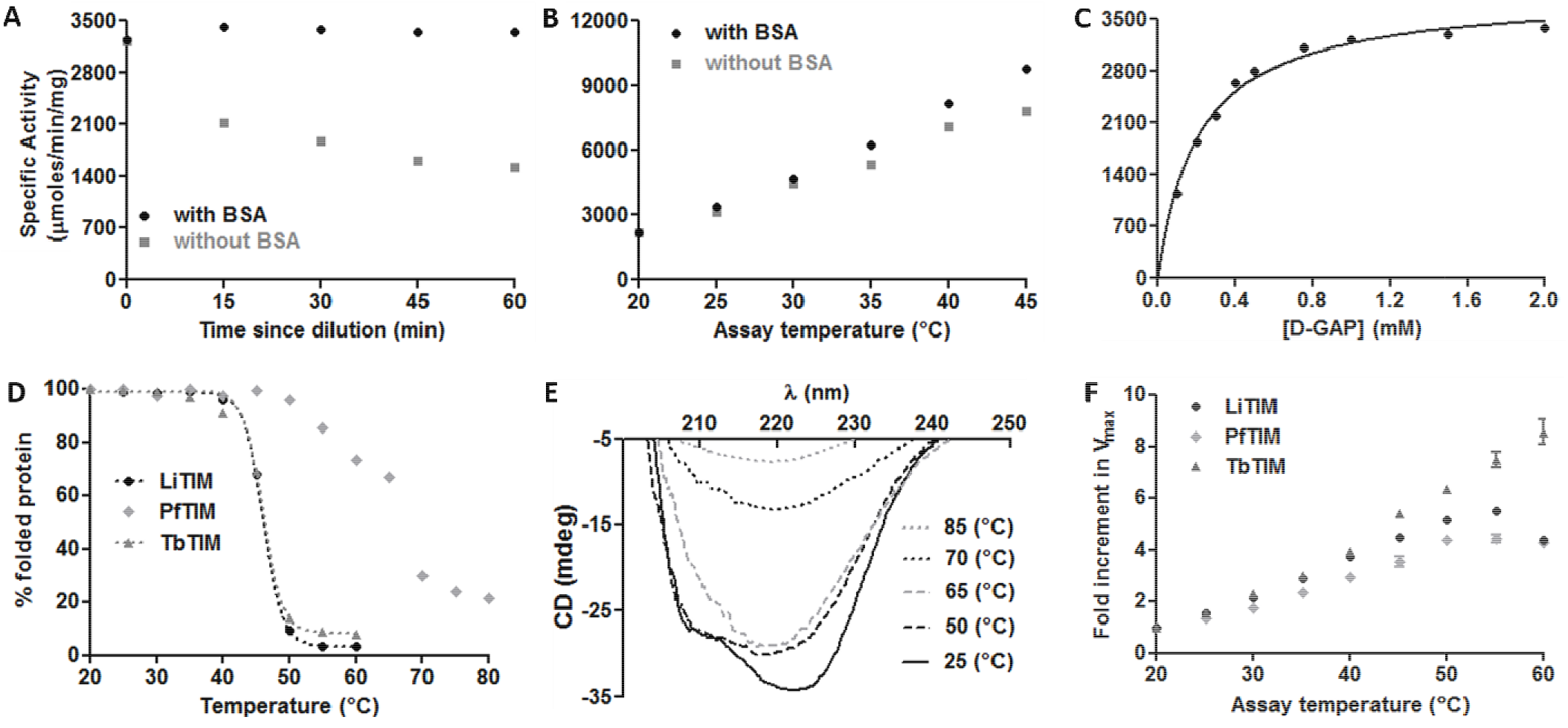
Biochemical characterization of LiTIM and comparison with TbTIM and PfTIM. The reduction in specific activity, upon (**A**) incubation of diluted enzyme at 4 °C (assay done at 25 °C) and (**B**) when the assay temperature was increased. The activity was rescued by adding BSA to enzyme solution and the assay mix (BSA alone showed no activity, data not shown). (**C**) The specific activity values obtained by substrate (D-Glyceraldehyde-3-phosphate) titration. (**D**) Thermal melting of LiTIM, PfTIM and TbTIM was performed by incubating the protein at the specified temperature and recording CD spectrum. Change in ellipticity, θ, (CD, mdeg) at 220 nm was used to calculate the % folded protein. LiTIM and TbTIM show sharp decline in the 220 nm ellipticity owing to rapid precipitation beyond 40 °C. The thermal melting profile of PTIM was more complex. (**E**) PfTIM showed different pre-melting spectral states prior to precipitation. (**F**) Temperature dependence of activity for LiTIM, PfTIM and TbTIM. (All experiments were repeated at least thrice and were reproducible. The graphs (**A**) (**B**) and (**C**) are one representative experiment while, in (**F**) data points are the means and standard deviation from data of three experiments. Assays reported in (**C**) and (**F**) were done with BSA in the reaction mix).

**Table 1.** Catalytic parameters for triosephosphate isomerase (25 °C) from three pathogenic organisms (*Leptospira interrogans, Plasmodium falciparum* and *Trypanosoma brucei*).

| Protein | $k_{\text{cat}} * 10^2 \text{ (s}^{-1}\text{)}$ | $K_m \text{ (mM)}$ | $k_{\text{cat}}/K_m * 10^2 \text{ (mM}^{-1}\text{s}^{-1}\text{)}$ |
| --- | --- | --- | --- |
| *LiTIM* | 17.41 | 0.21 | 82.9 |
| *PfTIM* | 5.02 | 0.9 | 5.58 |
| *TbTIM* | 9.45 | 0.17 | 55.6 |

### Temperature dependence of stability

Thermal unfolding profiles of *Li*TIM, *Tb*TIM and *Pf*TIM were compared by following the change in the far-UV CD spectra of the three proteins (Figure 2D). Both *Li*TIM (*T*_m_ of 46.5 °C) and *Tb*TIM (*T*_m_ of 46.1 °C) showed substantial precipitation between 40-50 °C, while the CD spectral band shapes remained similar till 50 °C. *Pf*TIM exhibited a complex thermal unfolding pattern and different band shapes were observed prior to precipitation (Figure 2D and E). With rising temperature, the first change observed was the loss of intensity around 220 nm (likely due to change in the environment of the Trp11 side chain present near the dimer interface [48]) followed by the loss of the helix signature peak around 208 nm resulting in a β-sheet dominated spectrum. A plausible explanation for such an observation could be the higher stability of the *Pf*TIM β-barrel core compared to *Li*TIM and *Tb*TIM. Indeed, later examination of the protein sequences of *Li*TIM, *Pf*TIM and *Tb*TIM and their crystal structures reveal pertinent differences (Figure S2 and S3). Compared to both *Li*TIM and *Tb*TIM, the inner core of *Pf*TIM is more tightly packed with bulky hydrophobic or aromatic amino acids pointing into the centre of the barrel (Figure S3A). Additional ionic interactions and aromatic interactions across the barrel core presumably contribute to the increased stability of *Pf*TIM barrel (Figure S3A and B). Closer examination of the β-strand registry revealed that while in *Pf*TIM there are four backbone-backbone hydrogen bonds between strand-4 and strand-5, two of the terminal hydrogen bonds are either very weak or absent in *Tb*TIM and *Li*TIM (Figure S3C). These structural differences between the three TIMs rationalise the higher temperature stability of *Pf*TIM in comparison to *Li*TIM and *Tb*TIM.

### Chaotrope induced denaturation

Change in intrinsic protein fluorescence intensity and emission wavelength maxima (λ_max_em) were followed as a function of increasing chaotrope (urea or guanidine hydrochloride) concentration and the data was fitted to the standard two state melting equations (Graphpad Prism; Figure 3). The protein was incubated with increasing urea or GdmCl concentrations at room temperature for 1 hr to allow equilibrium unfolding. There was a gradual decrease in fluorescence intensity and shift of the λ_max_em as the chaotrope concentration was increased. The apparent mid-transition concentrations, C_m_, for urea and GdmCl were 2.6 M and 0.8 M, respectively (Figure 3). The λ_max_em shows substantial red shift from 324 to 348 nm between the native and the fully unfolded protein (characteristic of a change in Trp environment from buried to solvated). In comparison, the reported apparent C_m_ for GdmCl induced unfolding of *Tb*TIM is ~1.6 M [55] and that for *Pf*TIM is ~1.5 M. Also, *Pf*TIM is known to retain some secondary and tertiary structure even at 8 M urea [56].

**Figure 3.**
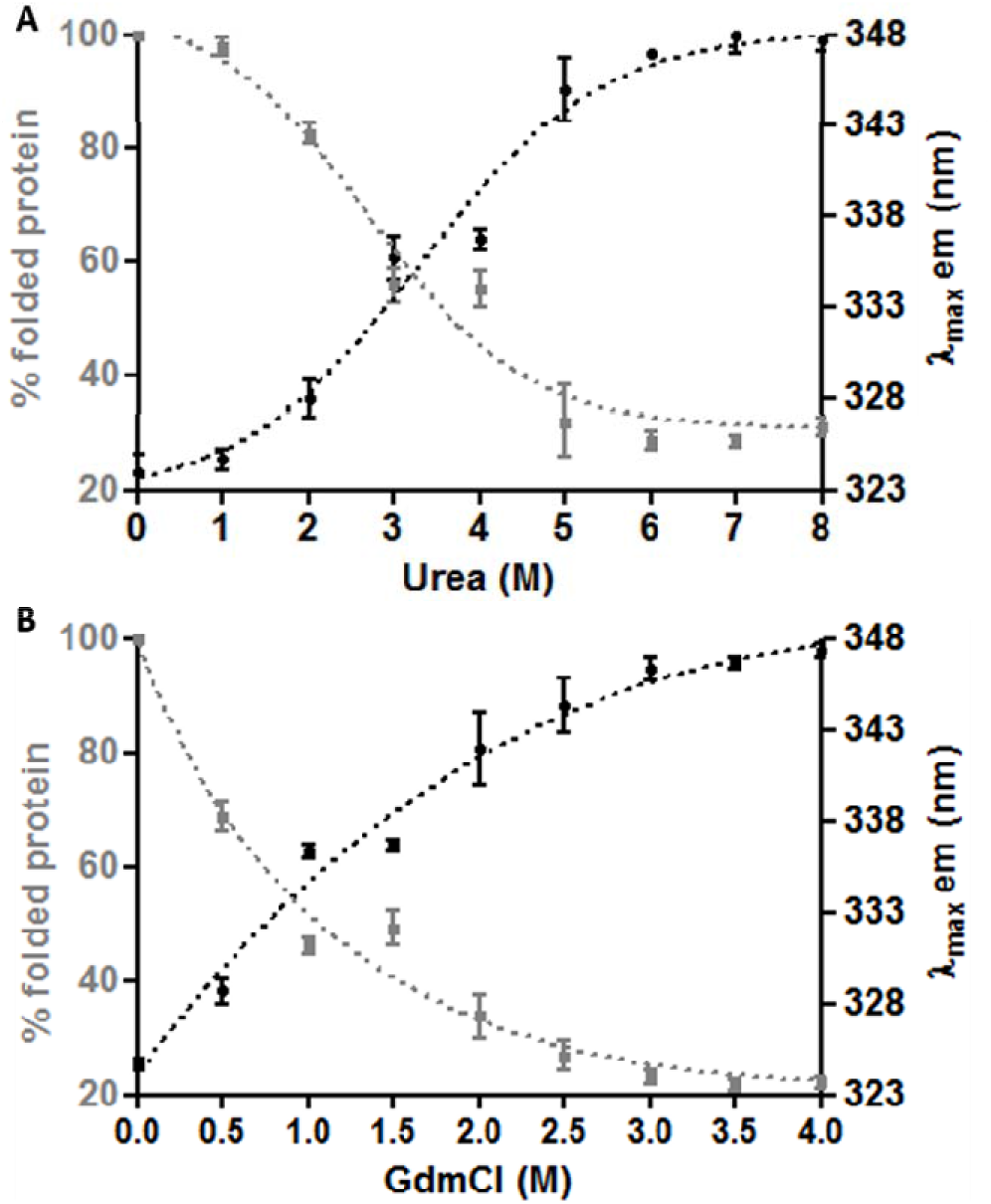
Chaotrope (urea and guanidine hydrochloride) induced denaturation of LiTIM. (**A**) and (**B**) show urea and guanidium hydrochloride (GdmCl) induced denaturation of LiTIM protein. The apparent denaturation midpoint concentrations, C_m_, were, ~ 2.6 M and 0.8M for urea and GdmCl, respectively, at room temperature. The change in the maximum emission wavelength (λ_max_ em = 324 nm) and the decrease in the fluorescence intensity at the maximum emission wavelenth of the native protein was monitored using excitation wavelength as 280 nm. (Standard deviation was obtained from data of three independent experiments).

### Temperature dependence of activity

Temperature and chaotrope induced unfolding captures the effect of sequence variability on the structural stability of proteins. Both, higher protein concentration, and longer incubation times (to reach equilibrium), contribute additively to the unfolding and precipitation. Equilibration of an enzyme at a higher temperature before determining its activity at a specific ambient temperature primarily reflects the effects of irreversible global structural perturbations. While rise in temperature may contribute to the rate enhancement by easing the essential dynamics of the protein [57–59], global melting and precipitation reduce the soluble active population of the enzyme. To delineate the contribution of these effects, we measured the Initial rate (or zero point rate) of *Li*TIM, *Tb*TIM and *Pf*TIM at low protein concentration. Studying a single enzyme from three different organisms has the advantage that only the non-catalytic residues vary, while the overall structure, oligomeric state, catalytic residues and their geometry are identical. We compared the effect of temperature on the specific activity profile of *Li*TIM, *Tb*TIM and *Pf*TIM under the same assay conditions and a saturating substrate concentration of 2mM (Figure 2F). Measurements at 25 °C showed that *Pf*TIM (*K*_m_= 0.87 mM) had lower substrate affinity compared to *Li*TIM (*K*_m_= 0.21 mM) and *Tb*TIM (*K*_m_= 0.17 mM). Variation of *K*_m_ of *Pf*TIM was determined in the assay temperature range and was found to remain the same (data not shown). Assays could be performed reliably till 60 °C, without any effect of the coupling enzyme denaturation (data not shown). A linear rise in specific activity was observed with rise in temperature from 20 °C to 50 °C for all the three enzymes. *Pf*TIM activity saturates at 50 °C, *Li*TIM shows highest activity at 55 °C and *Tb*TIM shows a rise till 60 °C. Clearly, temperature induced change of initial (or zero point) rate of reaction do not reflect the overall structural stability of the three proteins. Before the effects of overall unfolding and precipitation can set in, local conformational changes, presumably proximal to the active site, appear to contribute to the observed temperature dependence of activity.

### Crystallization, structure solution and analysis

*Li*TIM crystals were obtained in the absence of any ligand (Figure 1D). To obtain the DHAP bound crystal form, pre-formed crystals were soaked in crystallization cocktail containing DHAP (Table S2). Both the structures were refined to reasonable *R*_work_ and *R*_free_ (Tables 2 and 3). The native crystals (crystal 1) corresponded to the *P*3_1_21 crystal form, with a monomer in the asymmetric unit and no ligand bound to the active site (*apo* form). The crystals after soaking with substrate (crystal 2) corresponded to the *P*3_2_21 crystal form with a dimer in the asymmetric unit. Crystal-2 also showed blobs of residual electron density at the active site of both the subunits, which could not be accounted for in terms of water or ethylene glycol (the cryoprotectant) molecules. After careful inspection of the electron density map in crystal 2, the substrate DHAP was modelled with partial occupancy of 0.6 and 0.8 and water and sulfate with 0.4 and 0.2 occupancy in subunits A and B, respectively, Figure 4. Loop-6 was found in an *open* conformation in the *apo* form and *closed* conformation in the *holo* form of the enzyme (Figure 6 and Table S3). The 8 β-strands constituting the core of the classical (α/β)_8_ barrel fold are depicted in Figures S4A and S4B. The dimer interface is constituted by β→α loop-1 (residues 12-18), loop-2 (residues 45-51), loop-3 (residues 66-88), loop-4 (104-106) and helix-4 (97-103) of both the subunits burying a large surface area (~1750 Å^2^). The presence of a pair symmetry related aromatic clusters, Y69, H70, F76, F110 of one subunit and F103 and of the adjacent subunit, at the dimer interface would substantially stabilize the dimer interface (Figure S4C). The 2-fold symmetry axis that generates the native dimer passes through the dimer interface. Superposition of the *Li*TIM dimer with TIM dimers from other organisms revealed that the orientation of the monomeric units is variable in TIMs from different sources (for TIMs from any two organisms, the angle between the dimer 2-folds was found to vary between 1-13° for, data not shown). The 2-fold dimer symmetry axes of *Li*TIM and yeast TIM (PDB ID: 1NEY, the only other substrate bound TIM structure) make 11.5° angle (Figure S1).

**Table 2.**
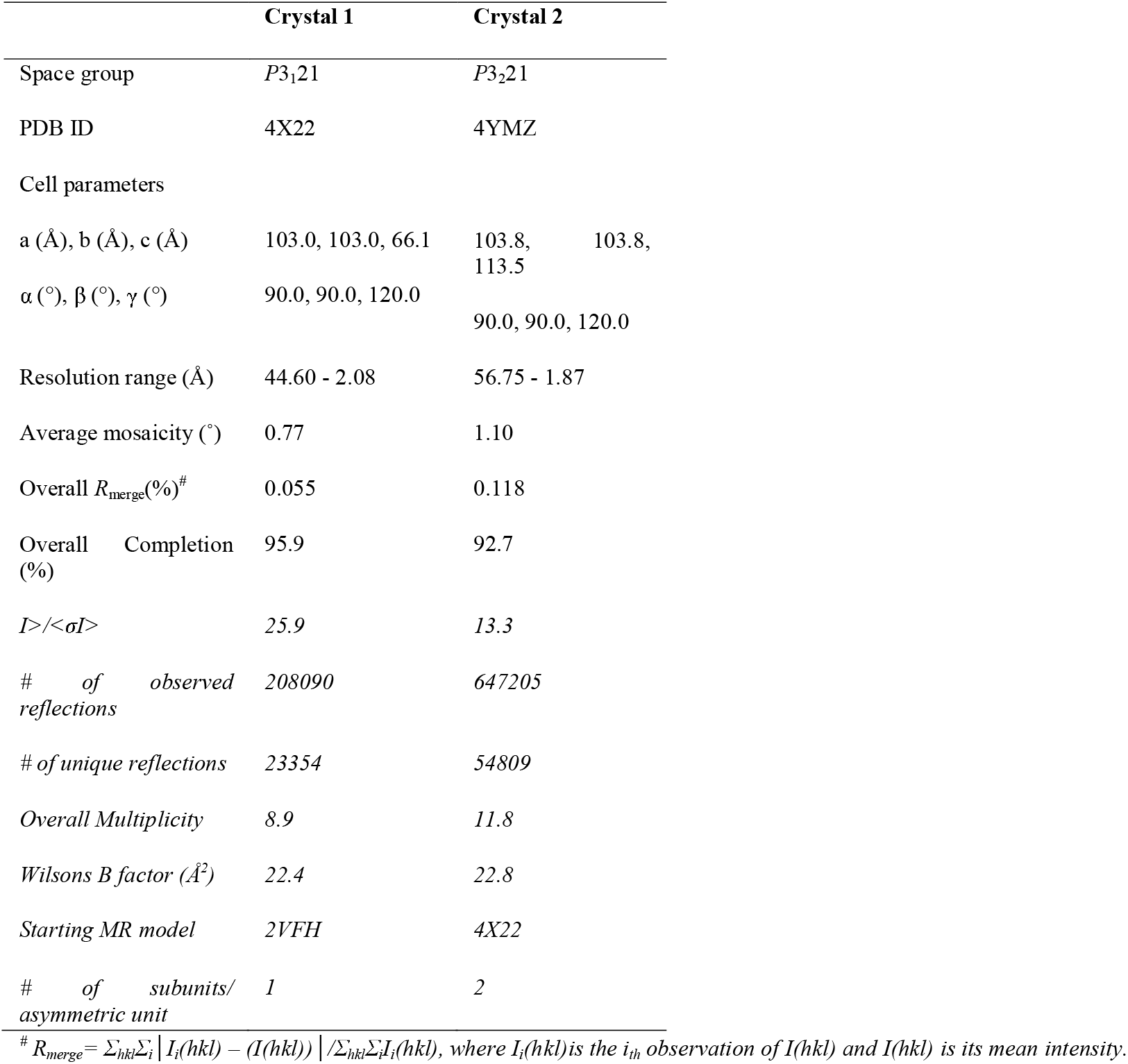
Data collection and processing statistics for *Li*TIM crystal 1 (*apo* form) and crystal 2 (*holo* form).

**Table 3.** Data refinement statistics for LiTIM crystals.

|  | *Crystal 1* | *Crystal 2* |
| --- | --- | --- |
| *Number of used reflections* | 22141 | 52007 |
| *% observed* | 95.1 | 92.8 |
| *R<sub>work</sub>(%)* | 15.4 | 17.1 |
| *R<sub>free</sub>(%)* | 19.1 | 21.1 |
| *FOM* | 89.7 | 88.2 |
| *Model quality:* |  |  |
| *Number of protein atoms* | 1874 | 3739 |
| *Number of water molecules* | 221 | 308 |
| *Number of ligand atoms* | 24 | 90 |
| *Matthews coefficient* | 3.71 | 3.23 |
| *Average B-factor ($\text{\AA}^2$):* | 23.0 | 27.8 |
| *RMS deviation from ideal:* |  |  |
| *Bond length ($\text{\AA}$)* | 0.02 | 0.02 |
| *Bond angle ($^\circ$)* | 1.89 | 2.01 |
| *Chiral volume ($\text{\AA}^3$)* | 0.12 | 0.13 |
| *Ramachandran statistics:* |  |  |
| *Favored region (%)* | 97 | 97 |
| *Allowed region (%)* | 3 | 3 |
| *Disallowed region (%)* | 0 | 0 |

**Figure 4.**
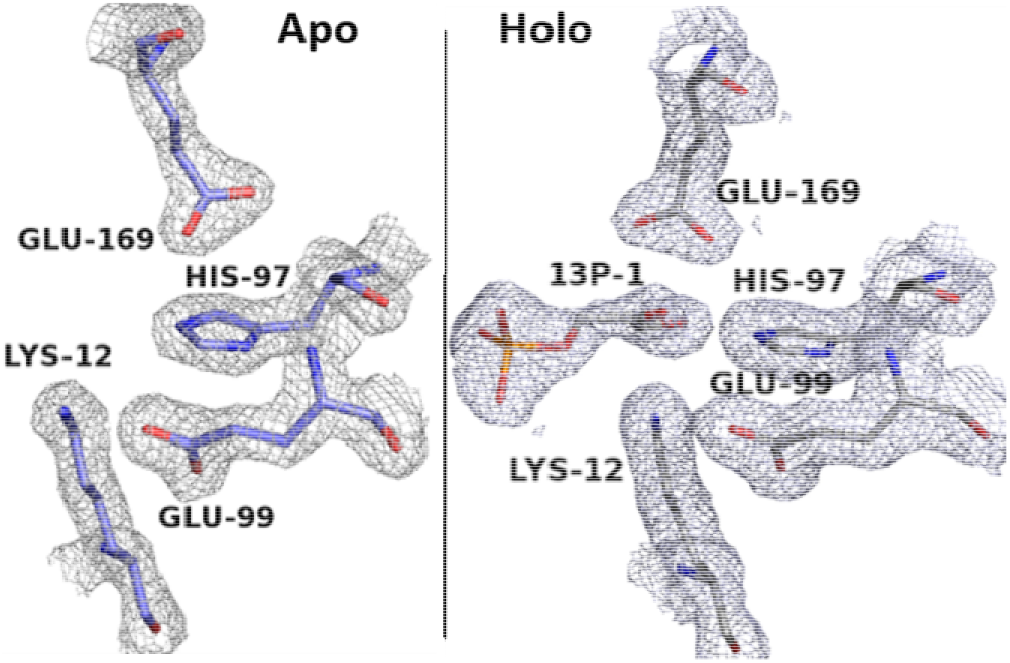
Active site residues and substrate (DHAP) electron density map in apo and holo forms of LiTIM. Electron density map (2Fo-Fc map contoured at 1σ) for the active site residues K12, H97, E99 and E169 in the apo (blue) and holo (grey) form structures of LiTIM. In the unmodelled blob of density at the active site of the DHAP soaked crystals, a dihydroxyacetone phosphate (13P-1) molecule was modelled.

### Active site organisation

Clear density was observed for the four catalytic residues§, K12, H97, E99 and E169 in both the *apo* and the *holo* forms (Figure 4). Comparison of the two states shows that out of these residues, only E169 (the catalytic base) shows an appreciable change in the backbone (φ changes from 104° to 130°; ψ changes from 116° to 98°) and side chain conformation (χ^1^ from −62° to −66°; χ^2^ from −151° to −179°; χ^3^ from 122° to 80°; and χ^4^ from −58° to −90°, Table S3) leading to the displacement of the carboxy terminus of the side chain by ~2.2 Å towards the substrate. Thus, the catalytic base E169 takes two distinct conformations, *swung out* and *swung in*, in the *apo* and the *holo* state of the enzyme, respectively (Table S3, Figures 4 and 8). The *swung out* conformation is stabilized by interaction with the imidazole ring of His97 and backbone −NH and side chain −OH of Ser98. The *swung in* conformation is attained by movement of E165 side chain away from the main body of the protein and into the active site cavity. In this conformation, the E169 side chain carboxy oxygens are at ~2.9 Å and 3.0 Å, and ~3.1 Å and 3.4 Å distance from the C1 and C2 of the substrate DHAP, respectively. The carboxy oxygen which is nearer to the substrate is also at a distance of ~3.3 Å from the back bone carbonyl −CO of the Leu238. This opens the possibility of a hydrogen bond upon protonation of this oxygen. Since the side chain of the catalytic base, E169, will be protonated after proton abstraction from the substrate, a hydrogen bond between the carboxylate oxygen and the L238 backbone carbonyl might contribute to the transition state stabilization. The structural aspects of this carboxy oxygen make it the more likely candidate out of the two side chain carboxy oxygens of E169. The other carboxy oxygen (which is relatively away from the substrate) remains at a distance of ~3.2 Å from the H97 side chain −N^ε2^ (believed to be protonated throughout the catalytic cycle). Also, after coming in proximity to the substrate the E169 side chain makes van der Waals contact with the side chains of the residues Ile174 (loop-6) and Leu236. These interactions involving the E169 carboxylate side chain will contribute to the shift of pKa observed for this residue [60–62] and thus be crucial for catalysis. One may note that the E169D (*Li*TIM numbering) mutant, studied by Knowles and co-workers, will lack all these interactions and may 1) perturb the desired pKa for the catalytic base, 2) increase the flexibility of the side chain involved in proton abstraction, 3) affect the transition state stabilization, and 4) increase the distance of the carboxy oxygen from the substrate, thus rationalizing the observed 1500 fold drop in the *k*_cat_ of this mutant [63].

The active site residue, Lys12 takes up a steriochemically disallowed backbone conformation (φ~ 44°, ψ~ −137°), as has been noted as a conserved feature in all the known functional TIM structures [64] (in monomeric variants of TIM this conformation is perturbed [65]). There are four apparent interactions that seem to be important for K12 conformation. 1) The side chain-backbone hydrogen bond between Asn10 and Lys12 (O ^δ^_i_… backbone NH _i+2_), and 2) backbone-backbone hydrogen bond between Trp11 and Asn14 (CO_i_… NH_i+3_) (Figure 5A). Contrary to our observation for the *Li*TIM structure, the backbone −CO of Trp11 has previously been observed in other TIMs (eg. *Pf*TIM: 1O5X, *Sc*TIM: 1NEY, *Tb*TIM: 6TIM) to be stabilized by backbone-side chain hydrogen bond between Trp11 and Asn14/His14/Tyr14 (CO_i_… N ^δ^ _i+3_) instead of backbone-backbone interaction [64] (Figure 5A). 3) The Lys12 backbone −CO makes a hydrogen bond with the side chain −OH of Ser45 and −NH_2_ of the amide side chain of Gln65. Residues 45 and 65, both being a part of dimer interface and making important inter-subunit interactions, emphasizing the importance of dimerization for TIM activity. The network of hydrogen bonds provides conformational stability to the peptide stretch from residues 10 to 14 and positions Lys12 into the active site. 4) K12 has a fully stretched side chain conformation (side chain dihedral angles ~180°) and the side chain NH_3_^+^ is within hydrogen bond distance from the bridging oxygen and the C_2_ carbonyl oxygen of the substrate on one side and the side chain carboxyl −CO of E99 on the other side.

**Figure 5.**
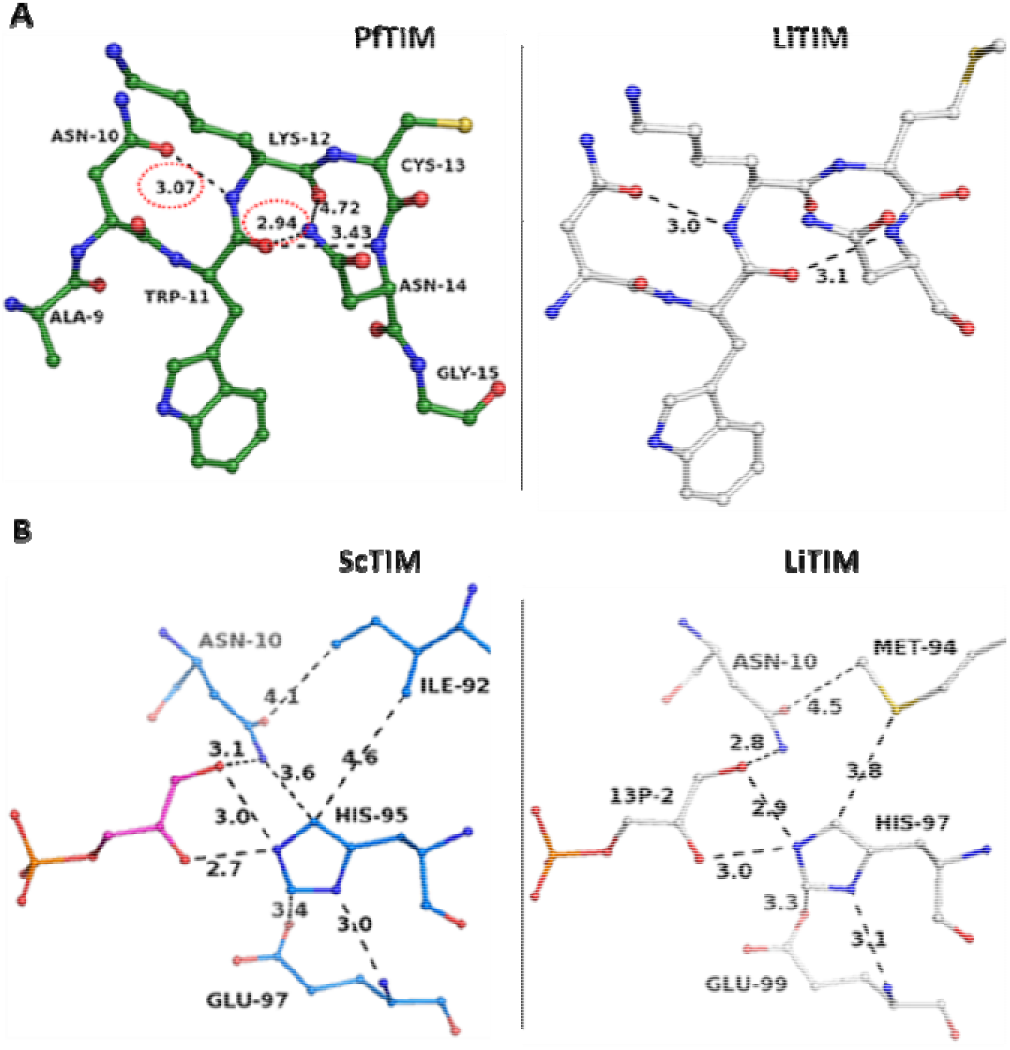
Subtle differences in the peripheral interactions of LiTIM active site residues compared to previously reported structures. (**A**) The (n-2) to (n+2) stretch of peptide around the active site residue Lys12 (LiTIM sequence numbering) showing how its strained backbone conformation (φ~ 50, ψ~ −120) is maintained by side chain-backbone hydrogen bonds between K12 … N10 (C_10_ ring, NH_i_… O^δ^_i-2_) on one side and either backbone-side chain hydrogen bond between N14… W11 (C_14_ ring, N^δ^_i_… CO_i-3_, as seen in PfTIM) or backbone-backbone hydrogen bond between N14… W11 (C_10_ ring, NH_i_… CO_i-3_, as seen in LiTIM). (**B**) N10 and the neutral imidazole form of H97 (LiTIM sequence numbering) neutralize the developing negative charge on the reaction intermediate. Unlike most other studied TIMs with a hydrophobic residue at position (i-3) from the active site His (eg. ScTIM), Met at position 94 in LiTIM or Glu at 124 in CiTIM may affect the pKa of the active site His (Figure S3B).

The other two active site residues, His97 and Glu99, are a part of a short helix spanning residues 97 to 104. Dimer interface interactions appear important for conformational rigidity of H97 as well as E99. The H97 imidazole ring lies parallel to the plane of branched side chain of T77 of the adjacent subunit (distance ~3.6Å) and the E99 side chain carboxy group is held in position by hydrogen bonds with T77 backbone −NH_2_ and side chain −OH (Figure S5A). The neutral imidazole ring of His97, can arguably adopt two alternative rotameric conformations- 1) N^δ1^ pointing towards the E99 backbone −NH (making an −NH… N hydrogen bond) and N^ε2^ making H-bond with E169 side chain; 2) N^δ1^ pointing towards residue M94 S^δ^ side chain and N^ε2^ pointing in the direction of E99 side chain carboxylic acid (Figure 5B). While there has been no experimental observation of the latter rotamer state, it might be an intermediate during catalysis as per an alternative TIM mechanism proposed by Samanta *et al*. [66]. This alternate mechanism would involve His97 as a proton acceptor [66], rather than as a proton donor as envisioned in the classical TIM mechanism [24]. Closer examination of H97 reveals that its side chain comes in close contact with the residue 94 (Met in *Li*TIM) side chain (S^δ^-N^δ^ distance- 3.6 Å) and presents the possibility of a sulphur-containing H-bond (as defined in [67, 68]) in the latter rotameric state of the His97 side chain (Figure 5B). Unlike *Li*TIM, in all the other reported TIM structures, this residue is a branched chain hydrophobic residue, Ile/Leu/Val, Figure 5B and Figure S2. A rare example of Glu at position 94 (*Li*TIM numbering) is in *Coccidiodes immitis* (PDB ID: 3S6D; Figures S2 and S5B).

### Loop movements and active site reorganisation

The active site loop-6 showed clear electron density conforming to the *open* and *closed* states in the *apo* and the *holo* form structures, respectively (Figure 6A, Figure 7 and Figure S6). The difference in the backbone dihedral angles of the stretch from residues 168-182 between the *apo* and the *holo* forms clearly shows the transition of loop-6 from *open* to *close* conformation (Table S3). In the *open* conformation, the residues at the tip of the loop-6 have no stabilizing interactions and thus possess high B factors. In comparison, upon closure, new interactions stabilize these residues, thus reducing the B factor of the loop-6 tip residues (Table S3; Figure S4). While the loop-6 *open* conformation allows entry of the substrate into the catalytic pocket (Figure 7A), loop closure ensures right positioning, tighter binding of the substrate and dehydration of the environment of the substrate (Figure 7C). In the *closed* state, residue Gly178 backbone −NH makes H-bond with the phosphate anion. In addition, the change of loop-6 conformation is accompanied by a corresponding change in the loop-7 conformation (Figure 6B), allowing Ser217 to make H-bond with the phosphate oxygen of the substrate (Figure 7C).

**Figure 6.**
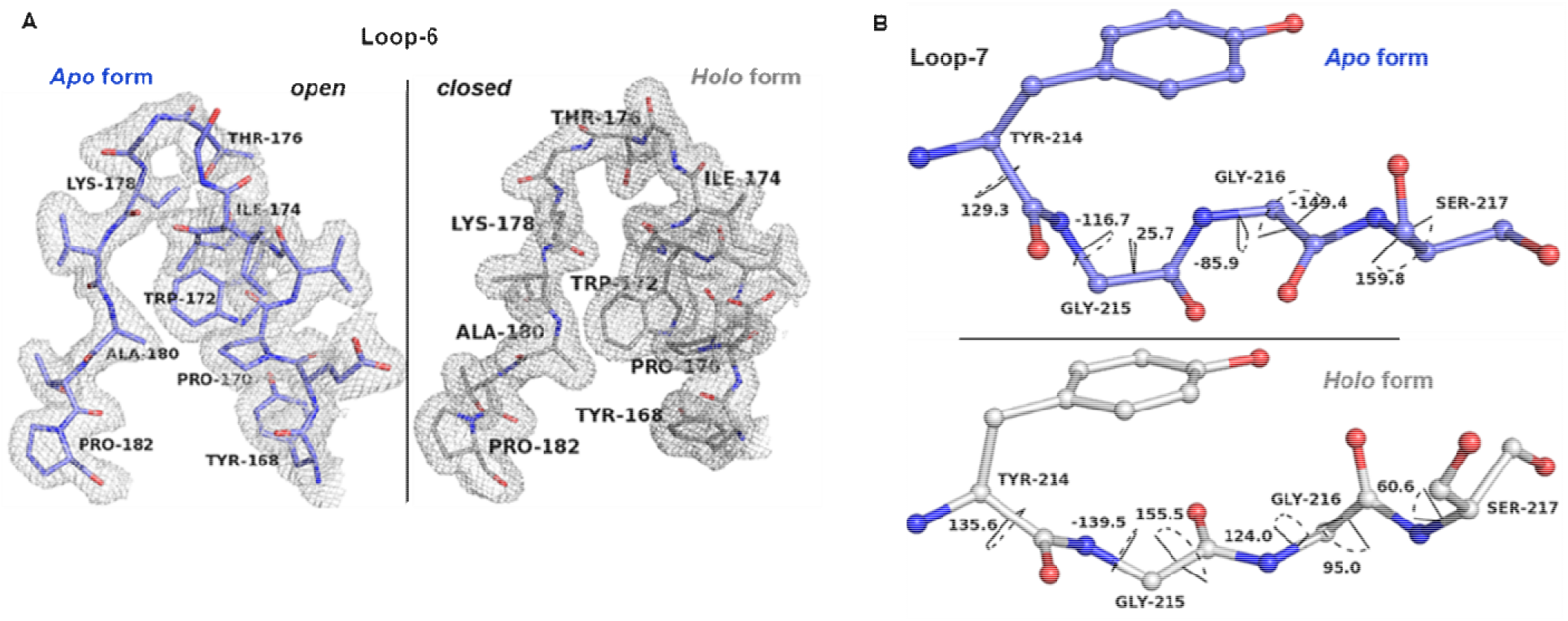
Loop-6 and loop-7 conformations in the apo and the holo states. (**A**) The electron density maps (2Fo-Fc contoured at 1σ) for loop-6 stretch from residue 168-182 in apo and holo (DHAP bound) form of LiTIM. There was clear difference between the two forms and while open conformation was observed for the apo form, closed conformation was observed for the holo form. (**B**) Transition in the backbone conformation of the loop-7 residues upon substrate binding-Gly215-Gly216 peptide unit flips 120°.

**Figure 7.**
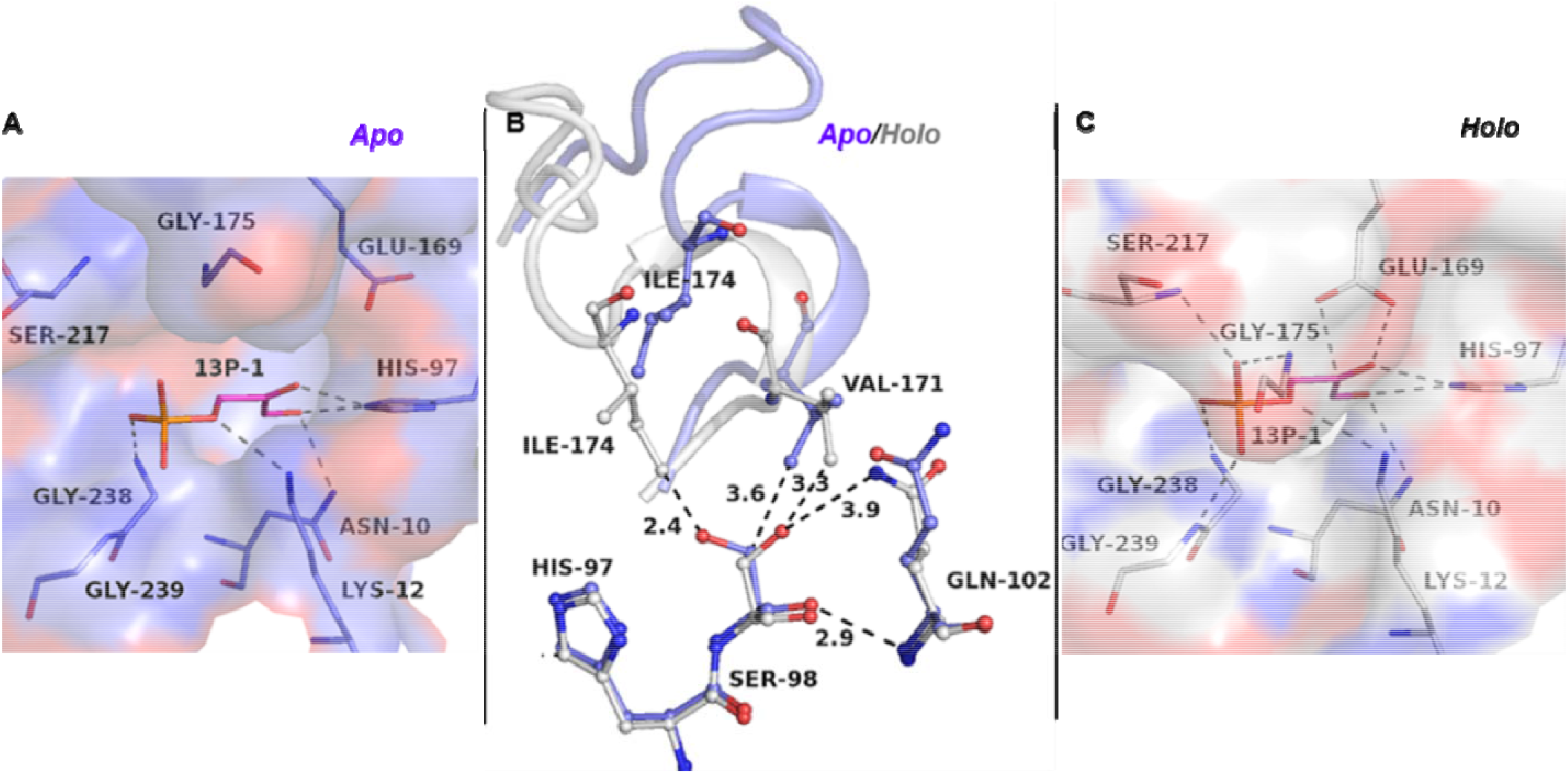
Reorganisation of TIM active site upon substrate binding and loop-6 closure. Transition of LiTIM active site from (**A**) the apo (blue) to (**C**) the holo (grey) state (substrate DHAP has been shown in both apo and holo forms to highlight the entry tunnel closure in the holo form). Loop-6 (residues 171-181) and loop-7 (residues 216-219) undergo conformational change to grip the phosphate end of the substrate and sequester the substrate from bulk solvent by closing over the active site pocket and place Glu169 proximate to the substrate. (**C**) Overlay showing loop-6 open and closed states. In the loop-6 closed state, the loop-6 hinge residue V171 and loop-6 tip residue I174 have a steric clash with apo state conformation of Ser98 side chain pushing it out of active site. Upon substrate binding and loop-6 closure, Ser98 flips from ‘gauche’ to ‘trans’ and Gln102 stabilises the ‘trans’ conformation by making a H-bond with Ser98 side chain hydroxyl.

Another subtle conformational change in the active site pocket, with plausible mechanistic implications, is that of residue Ser98 from *gauche* (*χ*^1^~ 45°) in the *apo* form to *trans* trans (χ^1^ 180°) in the *holo* form. Overlay of the *apo* and the *holo* form structures (Figure 7B) shows that the loop-6 tip residue Ile174 comes into unfavourably close contact with residue 98 side chain. This strain is relieved upon a change of conformation of the Ser98 side chain to the *trans* form. In the *gauche* conformation, residue 98 side chain stabilizes the E169 *swung out* conformation and upon ligand binding, while E169 moves into the catalytic pocket, Ser98 moves out of the catalytic pocket, interaction with Gln102 side chain stabilising this conformation (Figure 7B).

### Substrate binding

While TIM structures from 34 different organisms are available in the *apo* form and several inhibitor-bound structures are also available, a substrate bound structure is still a rarity. The only DHAP bound structure reported thus far is that of yeast TIM (PDB ID: 1NEY) [69]. In the substrate bound *holo* form we observed partial occupancy for the substrate. This seems plausible, given that we performed the crystal soaking experiment for limited time to avoid crystal cracking and dissolution. Binding of the substrate in the TIM active site presents a case where both preorganization and re-organization of the active site contribute to ligand binding and catalysis (Figures 6, 7 and 8). While some of the residues lining the active site (N10, K12 and H97) provide the platform for binding of the approaching sugar end of the substrate (Figure 7A and Figure 8B), it is the loop-6 and the loop-7 conformational changes that ensure tight anchoring of the phosphate end (Figures 7C and 8C) and positioning of the catalytic base E169, making the enzyme catalytically competent (Figure 7 and Figure 8A). The N10 side chain amide nitrogen (N10 N^δ^… DHAP O1 ~3Å) and K12 side chain terminal amine (K12 N^ζ^ … DHAP O2 ~ 3Å and K12 N^ζ^ … DHAP O3 ~ 3.2Å) make hydrogen bonds with the sugar moiety of the substrate. This would energetically favour binding of DHAP in an extended conformation rendering its carbon backbone exposed for proton abstraction by E169. H97 imidazole N^ε2^ is equidistant from both O1 and O2 (~3Å) and is likely to have strong interaction with the substrate as well as the ensuing negative charge on the enediol/ate intermediate (Figure 8B).

**Figure 8.**
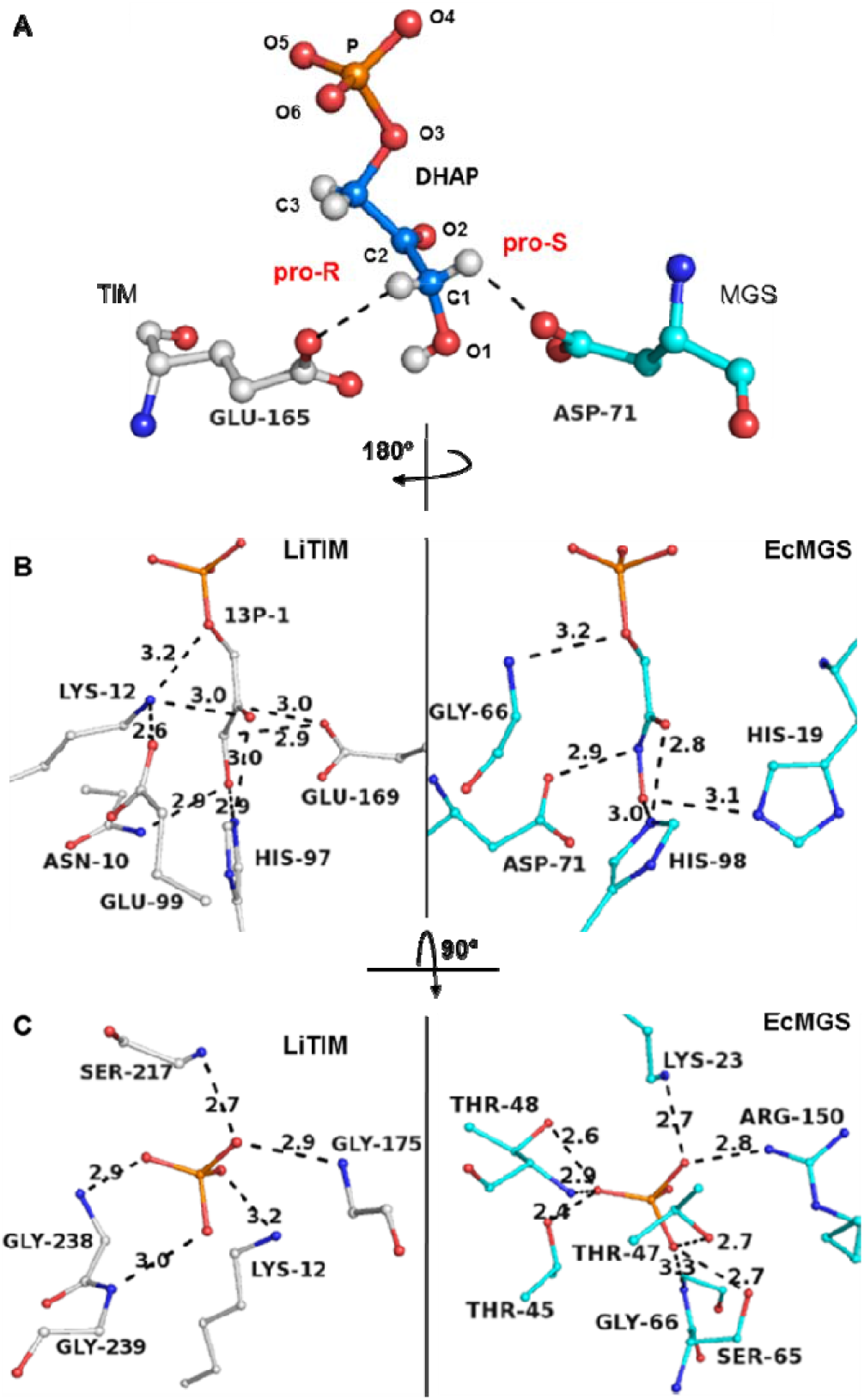
Ligand binding in triosephosphate isomerase versus methylglyoxal synthase. Three different orientations of dihydroxyacetone phosphate, DHAP/ phosphoglycolohydroxamic acid, PGH are shown for comparison of ligand binding mode in triosephosphate isomerase (TIM, grey) versus methylglyoxal synthase (MGS, cyan). (**A**) Overlay of yeast TIM (PDB ID: 1NEY) and E. coli MGS obtained by superposing the phosphate group of bound DHAP and PGH alone and moving the rest of the structure. The catalytic base (E165/D71) approaching the substrates from opposite directions ensure stereospecificity of the first proton abstraction. (**B**) and (**C**) Active site organization in TIM v/s MGS around the 3-C sugar backbone and phosphate group of the substrate (DHAP)/inhibitor (PGH), respectively. Orientations in (**B**) and (**C**) are related by 90° rotation along the C2-C3 bond axis. For clarity only the side chains of the residues and in (**C**) only the phosphate end of the substrate is shown. Left panel shows LiTIM, PDB ID: 4YMZ and right panel shows E. coli MGS, PDB ID: 1IK3. (**B**) In TIM, the catalytic base E169 is poised to abstract the pro-R proton while in MGS, Asp-71 can abstract the pro-S proton.

The small sugar phosphate, DHAP, is recognized by several enzymes in the cell and undergoes specific reactions in each case [70, 71]. Comparison of the modes of substrate binding, conformation of DHAP in the bound form and its interactions with the enzyme active site, in TIM and other DHAP bound enzyme active sites reveal functionally important differences (Figure 8, Figures S7 and S8, Tables S4 and S5). In TIM and MGS active sites, DHAP binds in an extended conformation with both, O1 and O2, pointing towards the same face of the binding pocket. Notably, the catalytic bases E169 (TIM) and D71 (MGS) approach from the *re* and *si* faces of the substrate (with respect to the C2 carbonyl group), priming the respective enzymes for stereospecific proton abstraction from the C1 carbon *ie*., the pro-R proton or the pro-S proton in TIM and MGS, respectively (Figure 8A). Unlike this, in glycerol-3-phosphate dehydrogenase (GPDH) and fructose-1,6-bisphosphate aldolase/lyase (FBPA) active site binding favours a bent conformation of DHAP and the O1 and O2 point in different directions (Figures S7 and S8). The phosphate moiety of the substrate is anchored by hydrogen bonds (donor- acceptor distance 2.5≥ ≤4Å) involving either neutral (backbone −NH, Ser side chain −OH and Thr side chain −OH) or cationic amino acids (Lys, Arg and His side chains). In TIM, only the bridging phosphate oxygen (O3) shows H-bond interaction with a charged side chain of K12. The other five potential hydrogen bonds made by the phosphate oxygens are to neutral backbone −NH (Gly175, Ser217, Gly238, Gly 239) or to Ser side chain −OH. These help to anchor the phosphate oxygens, contributing to the substrate binding energy and stabilization of the *closed* conformation of the active site loop-6 closing the entry/ exit mouth of the TIM active site. In contrast to TIM, the phosphate group in DHAP bound to MGS has eight likely hydrogen bond interactions to neutral groups and 2 to charged side chains (Figure 8, Figure S8 and Table S6). The importance of these differences with regard to enzyme reaction rate and reaction specificity is elaborated in the discussion section.

## Discussion

The 3-carbon sugar phosphates- DHAP and GAP- participate in several metabolic pathways and undergo different reactions by being catalysed by different enzymes (Figure 9) [2, 13]. Thus, the equilibrium between GAP and DHAP is likely to dictate the metabolic flux in the direction of energy generation and amino acid metabolism or lipid metabolism [70, 72], glyoxalase cycle and inorganic phosphate [73, 74] and NAD production [75] (-via de novo biosynthesis of quinolinic acid in bacteria and archaea- [76–78]) pathways, determining the adaptability of organisms to their nutritional micro-environments. Generally, breakdown of the 6-carbon sugar, fructose-1,6-bisphosphate, as a part of the glycolytic pathway is the primary source of DHAP and D-GAP. Leptospires are atypical organisms that rely on β-oxidation instead of glucose utilization via glycolysis [12, 21], and glycerol is believed to be the chief source of DHAP [13]. Given this, TIM catalysed isomerization of DHAP to D-GAP will serve as the entry point into pyruvate production followed by TCA cycle (Figure 9). We have reported here the cloning, purification and detailed biochemical characterization of the *Leptospira interrogans* TIM (*Li*TIM, Figure 1). Further, we have reported the high resolution X-ray structure of the *apo* and the substrate (DHAP) bound, *holo*, forms of *Li*TIM, paving way for future work to find a specific *Li*TIM inhibitor. Detailed structural examination of *Li*TIM and its comparison with other pathogenic TIMs (*Tb*TIM and *Pf*TIM) revealed specific features contributing to structural and functional adaptations of these enzymes, Figures 2 and 3. The human and *Li* TIMs have ~ 38% sequence identity, leaving 154 positions dissimilar in a 251 residues long monomer (Figure S2). Further examination of these differences may be useful in the search and design a *Li*TIM specific inhibitor.

**Figure 9.**
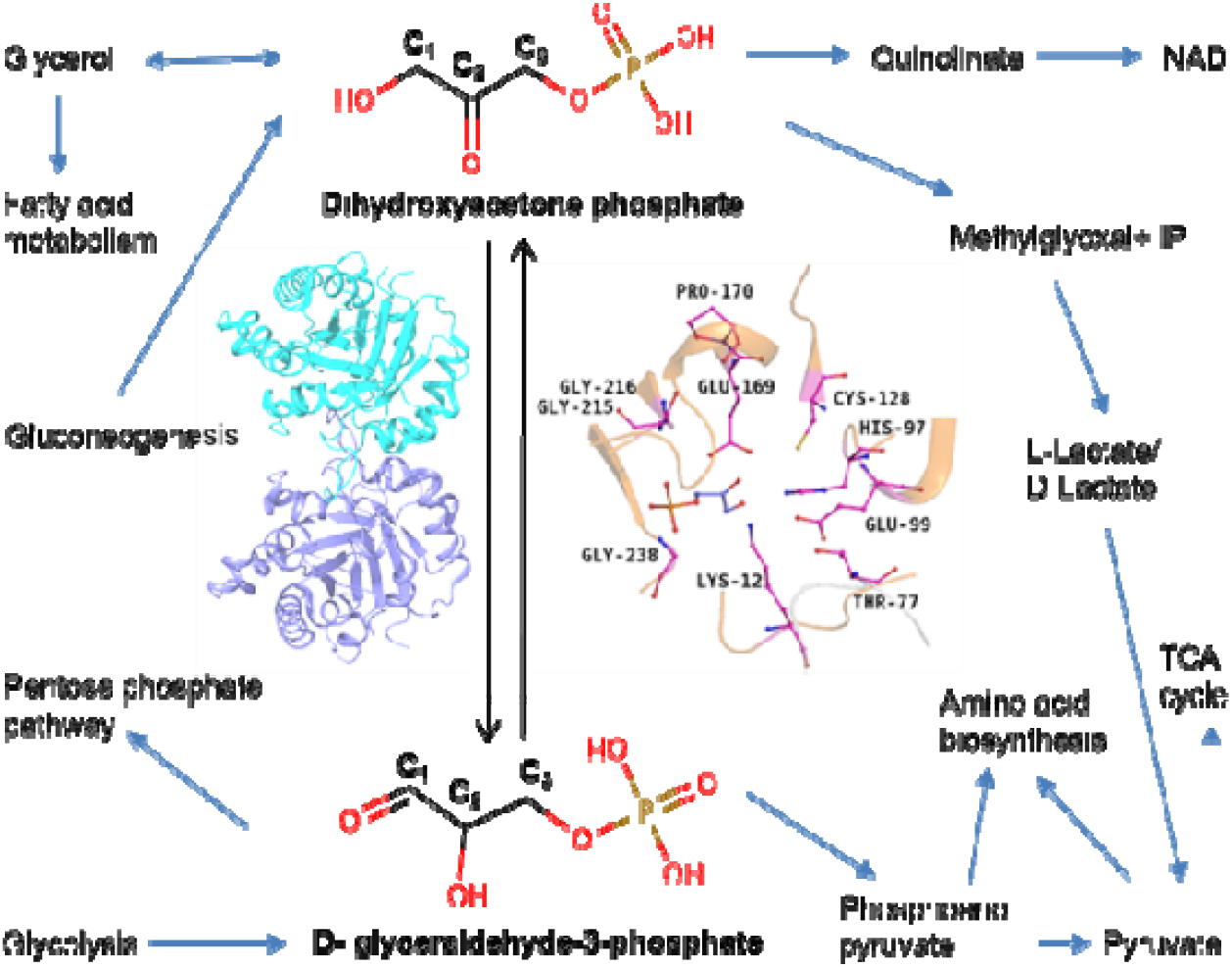
Place of TIM in a Leptospira metabolism map. The two sugar phosphates, dihydroxyacetone phosphate (DHAP) and D-glyceraldehyde-3-phosphate (D-GAP), participate in several essential metabolic pathways and triosephosphate isomerase catalysed interconversion will thus regulate the flux to different metabolic routes in Leptospires. LiTIM isomerises DHAP and D-GAP (The native dimer shown as cartoon representation and the conserved stretches that construct the active site of the enzymes are highlighted with the 100% conserved residues highlighted in magenta ball and stick.

### Role of replacements in enzymatic plasticity

Enzymes are protein scaffolds that bring reactive catalytic residues together in space, orient them in a specific geometry, thus imparting substrate and reaction specificity (Figure 9). This architecture is kept intact under the physicochemical environment of an organism. Protein sequences evolve under adaptive pressures but it is not straight forward to identify the contribution of variability of peripheral residues on the stability and activity of an enzyme. For the ubiquitous enzyme TIM, only 10 residues were found to be 100% conserved in a curated dataset of ~6300 sequences (unpublished), Figure 9. The remaining residues can be mutated to variable degrees still conserving the fold and function of the enzyme. Recently, the effect of the mutation E104D (a highly conserved position that has dimer destabilising effect and is related to human disease [79], numbering as human TIM) on the human TIM was compared with the same mutation done in the background of other pathogenic TIM sequences (TIM from *Giardia lamblia, Trypanosoma cruzi*, and *Typanosoma brucei*). The study showed that the mutation E104D affected the dimer stability to different extents in the four enzymes [80]. This observation suggests that non-conserved residues may make unique contribution to the global properties of a protein. For the three mesophilic TIMs examined in the present work, as many as 180 positions in a 250 residue full length protein are different (30% identity for the three sequences; Figure S2). We observed differential effect of temperature on the stability and activity of TIMs from the three human parasites *Leptospira interrogans, Trypanosoma brucei* and *Plasmodium falciparum* (Figures 2 and 3). While a more rigid inner β-barrel core (Figure S3) imparts greater thermal stability to *Pf*TIM, it does not seem to provide advantage with regard to the temperature dependence of activity (Figure 2). Local motions of residues H97, Ser98 and Glu169 [33, 66, 69] and loop-6/loop-7 dynamics is an integral part of TIM catalysis [81–83], Figure 7. Dimer integrity plays an essential part in maintaining active site geometry and thus may abate the unproductive effect of small thermal fluctuations. Notably, loop-6 opening-closing rates and catalytic rates are finely coordinated in TIM and its perturbation causes either loss of the enediol intermediate leading to methylglyoxal formation [81, 84]. The triad combination of residues at positions 98, 171 (loop-6 hinge) and 175 (loop-6 tip) connect the loop-6 movement to the precise positioning of E169 side chain upon substrate binding [33]. While the triad combination is SVI for *Li*TIM and *Tb*TIM, it is FLI for *Pf*TIM (Figure S2). Will the three TIMs show different temperature dependence of loop dynamics? The answer must await precise measurements. The interplay of increased productive dynamics as well as the temperature driven transition of the protein to unproductive configurations will both contribute to the change of activity with rise in temperature. The classical view of change of activity with temperature only considers denaturation being the cause of loss of activity at high temperature. Unlike this, the equilibrium model [85] attributes temperature dependence of activity to the effect of local motions on catalysis [86–89]. Our results can be rationalized by incorporating the contribution of sequence encoded local thermal fluctuations to enzyme catalysis.

### Achieving speed and specificity the TIM way!

The active site residues N10, K12, H97 are pre-organized to augment selective binding of the substrate conformation which allows efficient proton abstraction and intermediate stabilization. While this is an example of the celebrated lock and key mechanism for substrate binding and catalysis, active site re-organisation (Figure 7) *ie*., set of internal motions happening synchronous to substrate binding, is an integral part of TIM catalysis. Comparison of the *apo* and the *holo* structures of *Li*TIM reveals the following conformational changes in the active site pocket- 1) The catalytic base E169 moves from *swung out* to *swung in* conformation, coming in close contact with the substrate, 2) Loop-6 moves from *open* to *close* conformation accompanied by loop-7 conformational flip, 3) Ser98 moves from *gauche* to *trans* conformation (Figures 6 and 7). At this point, residues N10 and M94 need a special mention (Figure 5). While N10 interacts directly with the O1 of the substrate and the intermediate, and is found to be largely conserved in a unique sequence dataset (unpublished) of TIMs from diverse sources (Asn in ~97% of 6277 total sequences), Ser is a rare variation observed in 184 bacterial sequences and 5 eukaryotic sequences (Table S7). One such example is that of the fungus, *Coccidioedes immitis* TIM (PDB ID: 3S6D; *apo* form structure, Figure S5B). Residue 94 is predominantly a branched side chain, non-polar residues (Table S7) and is in van der Waals contact with the catalytic residue His97 side chain. *Li*TIM has a rare residue Met at this position and the S^δ^ makes a potential H-bond with His97 side chain (Figure 8, Figure S8). Such a H-bond will affect the side chain rotational freedom as well as affect the pKa of the imidazole ring [90, 91]. An even more drastic effect will ensue in cases where residue 94 is a Glu, as in *Coccidioedes immitis* TIM (Figure S5B). In considering the chemical reasonability of his proposed TIM mechanism, Knowles remarked that it “remains a puzzle” why the “enzyme uses imidazole rather than the much more electrophilic imidazolium as its acid component” [24]. In fact, he called it an “unexpected electrophile” and considered it a “seductive possibility that the proton transfers are mediated merely by the rotation of the appropriate side chain” [24]. More recently, this alternative to the classical TIM mechanism, which proceeds by rotation of the histidine ring and avoids involvement of H95 *imidazolate*, has been elaborated by Samanta *et al*, [66]. Sequence conservation analysis of TIM from diverse sources is currently in progress and will provide insights into evolution of catalytic mechanism and to identify functionally relevant rare variations in enzymes.

TIM catalysed isomerization is known to proceed via formation of an enediol intermediate. The partition of the enediol into elimination versus isomerization is 100 in the absence of TIM, indicative of the ease of phosphate elimination from the enediol intermediate [25, 32]. Juxtaposition of the active site of TIM and methylglyoxal synthase, MGS, (the enzyme that catalyses elimination reaction of DHAP) provide insight into how such remarkable reaction specificity is achieved by the two enzymes (Figure 8 and Figure S8). Comparison of the *holo* form of *Li*TIM with the phosphoglycolohydroxamic acid (PGH-enediol intermediate analogue) bound form of MGS shows molecular details of similarity and differences in the active sites of the two enzymes. 1) While E169 (TIM) is primed to abstract the C1 pro-R proton from DHAP, D71 approaches the substrate from the opposite direction and thus is suitable for abstracting the C1 pro-S proton. 2) After formation of the enediol intermediate, E169 can donate back the proton to the C2 carbon due to its proximity to both the carbons. In contrast, D71 is too far from C2 of the substrate. This also rationalizes the fact that D-GAP cannot act as a substrate for MGS. 3) Loop-6 closure in TIM ensures removal of bulk water from the active site and dehydration of the active site serves to quench the spontaneous elimination from the intermediate.

### Isomerization versus elimination- tale of the phosphate tail!

Phosphate binding energy is known to contribute ~12 kcal to the ΔG of isomerization by TIM [92]. The substrate that enters the TIM active site is only loosely bound by N10, K12 and H97, but loop-6/ loop-7 conformational changes provide a firm hold by positioning the hydrogen bond donors for interaction with the phosphate oxygens. Rate of isomerization drops by 10^5^ fold in the loop-6 deletion mutant [93]. Despite binding DHAP in similar configuration, TIM isomerizes it to D-GAP while MGS carries out elimination of the phosphate. Comparison of the interactions of phosphate in the two enzymes, TIM and MGS, clearly shows that the overall number of potential H-bond interactions as well as the number of polar/ charged interactions is greater in the case of MGS (Figure 8, Figure S8 and Table S6). The significance of this is in making phosphate a better leaving group and facilitating elimination reaction by MGS, while avoiding elimination in the TIM active site. In summary, the substrate bound structure of triosephosphate isomerase (*Li*TIM) from the zoonotic pathogen *Leptospira interrogans* provides insight into the structural imperatives for stereospecific proton rearrangement during TIM catalyzed isomerization and highlights the role of pre-organization and re-organization in achieving high reaction rate while maintaining reaction specificity. Kinetic, biophysical and structural investigations of the three mesophilic TIMs reveal that sequence variation at non-catalytic residues results in a tradeoff between temperature induced rate enhancement and unfolding. The variation of non-catalytic residues vicinal to H95, one of the catalytic residues, instigated a detailed study (unpublished) analyzing the sequence conservation pattern observed in all the available TIM sequence and investigation of minor structural changes in the TIM active site of different organisms that seem to have mechanistic implications.

## MATERIALS AND METHODS

### Cloning of the *Li*TIM gene

Triosephosphate isomerase gene (*tim* gene ID: LEP2GSC113_RS0110880) was amplified from five days old *Ellinghausen–McCullough–Johnson–Harris* (*EMJH*) culture of *<u>Leptospira interrogans</u>* (*Li*) serovar Icterohaemorrhagiae strain RGA (Regional Medical Research Centre, Indian Council for Medical Research, Port Blair, India; ATCC: 23581. Taxonomy ID: 1291351) by colony PCR (cell pellet from 30 μL of culture was resuspended in 60μL of autoclaved milliQ water followed by heating at 95°C for 20min. 15μL of the lysate thus obtained was used as template for 50μL PCR reaction mix). The PCR reaction was carried out using *Li*TIMFP and *Li*TIMRP (Table S1. Sigma-Aldrich Co, India) as the forward and the reverse primers, respectively, inserting NcoI and BamHI endonuclease sites at the 5 ’ of the start and 3 ’ of the stop codons of the gene, respectively. To circumvent the problem of frame-shift in the gene caused due to NcoI site at the 5’ end, two extra nucleotides were inserted downstream of the endonuclease site (underlined in the primer, Table S1). The extra codon (GCT) results in insertion of Ala immediately after the N-terminal Met of the protein. The PCR mixture contained, in a total volume of 25 μl: template DNA, 100 ng; mutagenic primer, 20 pmol; thermostable polymerase buffer (10X), 2.5 μl; dNTPs, 6 μl of a solution containing 2.5 mM of each dNTP; and polymerase 2 U. Following the hot-start PCR method, PR polymerase (Bangalore Genei, Bangalore, India) was added to the reaction mix after initial denaturation at 95°C for 5 mins. Subsequently, the steps, 1) 95°C for 15 sec, 2) 55°C for 20 sec and, 3) 72 °C for 90 sec repeated for 28 cycles followed by a final extension at 72°C for 5 min, to obtain the amplified PCR product which was then purified (Quiagen Gel purification kit) and digested with Nco1 and BamH1 restriction endonucleases (NEB, India) for inserting the gene into the pTrc99A expression vector. The ligated product was transformed in DH5α cells and plated on LB agar containing 100 μg/mL ampicillin. 10 colonies were screened for a positive clone (with *Li*TIM insertion) by carrying out restriction digestion with Nco1 and BamH1 enzymes on the plasmid obtained from the respective colonies and confirmation by primer specific PCR amplification, followed by confirmation with gene sequencing (final gene length 756 bp).

### Protein expression, purification and basic characterisation

*E. coli* expression strain AA200 (TIM null mutant) were transformed with pTrc99A containing the recombinant *Li*TIM WT gene and grown in LB for 8-10 hrs with 100 μg/ml ampicillin. The culture was used as 1% pre-inoculum in Terrific broth (TB) culture grown at 37 °C. After induction with 500 μM IPTG (OD_600nm_ 0.6-0.8), the culture was allowed to grow for another 16hrs at 30 °C. Cells were harvested by centrifugation (20 min, 6K rpm, 4°C), resuspended in lysis buffer containing 20 mM Tris-HCl (pH 8.0), 1 mM EDTA, 0.01 mM PMSF, 2 mM DTT and 10% glycerol and disrupted by sonication. After centrifugation (45 min, 12K rpm, 4 °C) and removal of the cell debris, the supernatant was fractionated with ammonium sulfate. The protein fraction containing TIM was precipitated between 35% and 65% ammonium sulfate saturation and the pellet obtained by centrifugation (45 min, 12K rpm, 4°C), was re-suspended in buffer A (20 mM Tris-HCl (pH 8.0), 2 mM DTT, and 10 % glycerol) and extensively dialyzed against buffer A at 4 °C. Further purification was done by anion exchange (Q-Sepharose, XK16 column, Amersham Biosciences) chromatography, with a linear gradient of 0-1 M NaCl and the fractions with purified protein were dialyzed against buffer A at 4 °C. Protein purity was checked on SDS-PAGE and LC-ESI/MS (Bruker Daltonics) was done to determine the correct mass of the protein. Electrospray ionization mass spectra were recorded on MaXis impact Q-TOF (Bruker Daltonics, Bremen, Germany) coupled to an Agilent 1200 series online HPLC. The molecular weight indicated (Figure 1G) is inclusive with an alanine inserted at the N-terminus of the protein. *Tb*TIM and *Pf*TIM were expressed and purified as described previously [33, 34].

### Spectroscopic characterization of the native protein

Circular dichroism spectrum was recorded on a JASCO-715 spectropolarimeter (JASCO technologies, Tokyo, Japan), with protein concentration of ~20 μM. Fluorescence emission spectra were recorded on a HITACHI F2500 spectrofluorimeter. Protein samples were excited at 295 nm and the emission spectra were recorded from 300 nm to 450 nm. Excitation and emission band passes were kept as 5 nm and 10 nm, respectively. All the spectra were corrected by subtracting the signal from buffer solution. Protein samples (final concentration ~5 μM) were prepared in 20 mM Tris HCl (pH 8).

### Analytical gel filtration

For determining the oligomeric state of *Li*TIM, analytical gel filtration chromatography was performed on a Superdex-75 (S75, GE Healthcare) 10 x 30 column. Gel filtration was done on a pre-equilibrated column with a flow rate of 0.2 ml/min, with Tris buffer (pH 8. 100 mM), 200 mM KCl and absorbance at 220 nm and 280 nm was recorded to monitor the elution of protein. Injection volume was 100 μl with protein concentration ~20 uM. To remove any precipitate or particulate impurity, protein was centrifuged at 14,000 rpm for 1 hr at 4 °C before injection. *Pf*TIM wild type protein was also run under the same conditions and used as an external control.

### Enzyme activity measurement

Enzyme activity was measured by a coupled enzyme assay method [35]. TIM catalysed conversion of D-GAP to DHAP was monitored in the presence of the coupling enzyme, α-glycerophosphate dehydrogenase (GPDH). The reaction mixture contained (final volume 500 μL) 100 mM TEA (pH 7.6), 5 mM EDTA, 0.5 mM NADH and 20 μg/m GPDH and GAP (for calculating the concentration it was assumed that the free acid solution contained D and L-GAP in equal concentration) to which TIM was added (5ul) to initiate the reaction. Substrate concentrations were varied from 0.2 mM-2 mM of D-GAP. The progress of the reaction was monitored at 25 °C by the decrease in absorbance of NADH at 340 nm (ε_340nm_= 6220 M^−1^cm^−1 [36]^). Upon addition of BSA solution (final concentration ~20μg/mL) to the diluted enzyme solution as well as to the reaction mix, the initial rates showed a linear dependence on the enzyme concentration in the range studied (data not shown). The values for the kinetic parameters (*K*_m_, *k*_cat_) were determined by fitting the initial velocity data to the Michaelis-Menten equation using Graphpad Prism (Version 5 for windows, Graphpad software, San Diego, California, USA, www.graphpad.com). The kinetic parameters for *Pf*TIM and *Tb*TIM were determined similarly. Protein concentration was determined by Bradford’s assay and also by measuring protein solution absorbance at 280 nm (theoretical value for ε_280nm_ *Li*TIM = 17,000, *Pf*TIM = 21,700, *Tb*TIM = 35,000 M^−1^cm^−1^, assuming all Cys reduced). At least three independent measurements of all assays were performed and the error was found to be within 5% of the mean.

### Thermal denaturation

For thermal melting studies, *Li*TIM, *Pf*TIM and *Tb*TIM (~20 μM final concentration in 20 mM Tris HCl, pH 8) were incubated at each temperature (20-85 °C with 5 °C temperature jumps) for 15 min for reaching equilibrium and three CD scans were performed at scan speed of 10 nm*min^−1^. CD measurements were performed on JASCO-715 spectropolarimeter (JASCO technologies, Tokyo, Japan) equipped with a thermostat cell holder controlled by a Peltier device. 1 mm path length cuvette, 2 nm band pass was used. The HT voltage remained in the permissible range throughout all the scans. Change in the CD ellipticity at 220 nm (θ in mdeg) was plotted as a function of temperature and the data for *Li*TIM and *Tb*TIM were fit to a monophasic transition model using Graphpad Prism (Version 5 for windows, Graphpad software, SanDiego, California, USA, www.graphpad.com).

### Chaotrope mediated denaturation

Equilibrium unfolding was carried out by incubating *Li*TIM protein (6*μ*M) with GdmCl (0-4 M) or urea (0-8 M) solution freshly prepared in 20 mM Tris-HCl (pH 8.0), for 1 hr. Emission spectra (310-450 nm) were recorded after exciting protein samples with 280 nm wavelength beam (5 nm band pass filter) on a Hitachi F2500 spectrofluorimeter. Each titration was repeated thrice and the respective buffer spectrum was subtracted. The mean and standard deviation of emission wavelength maxima (λ_max_ em) at each concentration and intensity at 324 nm were plotted to follow the change in the protein intrinsic fluorescence caused by protein unfolding. Data were fitted to monophasic transition model using Graphpad Prism (Version 5 for windows, Graphpad software, San Diego, California, USA, www.graphpad.com). Spectra were corrected by subtracting the buffer signal.

### Temperature dependence of activity

Reaction mix (500 μl) with TEA buffer, NADH, EDTA, BSA and required amount of water were incubated at the desired temperature for 5 mins and after adding 4mM D/L-GAP and coupling enzyme, reaction was initiated by addition of specific amount (5 μL of stock with 100 times higher concentration than the final concentration) of *Li*TIM (~3.3 ng/reaction) / *Pf*TIM (~6.0 ng /reaction) / *Tb*TIM (~5.7 ng / reaction) proteins. Protein solution was kept on ice and reaction was monitored immediately after addition of the enzyme. Kinetic measurements were done in the range of 20-60 °C with 5 °C interval.

### Crystallization, data collection, processing, structure solution and refinement

The purified *Li*TIM protein was concentrated to ~14-16 mg/ml and was centrifuged at 14,000 rpm for 1hr at 4 °C prior to setting up for crystallization. Diffraction quality crystals were obtained by the hanging drop vapour diffusion method (composition of the crystallization cocktail are given in (Table S2), at 23 °C; micro-crystals appeared within a week which were allowed to grow for a month to improve the crystal size and diffraction quality. Each hanging drop contained equal proportion of the protein and the crystallization cocktail, which was left to equilibrate with 450 μl of the crystallization cocktail as reservoir buffer. To obtain ligand bound *holo* form, crystals were incubated with ~280mM DHAP (solution made in the reservoir buffer) at 23 °C and were monitored for signs of cracks or crystal dissolution under a light microscope. For crystal-1 diffraction data were collected on the home source Rigaku rotating anode generator and MAR Research image plate detector system and 10 % ethylene glycol was used as cryoprotectant. After incubating with DHAP for around 25 min (crystal-2) (15% ethylene glycol as cryoprotectant), crystals were mounted on a cry loop flash frozen in liquid N_2_ and the X-ray diffraction data was collected on Beamline BM-14U at ESRF, Grenoble, France. Data were processed with iMOSFLM [37] and SCALA [38, 39], of the CCP4i suite of programs [40, 41] (Table 2). The data collected for crystal 1 and crystal 2 were processed in the trigonal space groups *P*3_1_21 and *P*3_2_21, respectively. Final structures were determined with the molecular replacement program PHASER 2.5.0 [42] of the CCP4 package. As the MR search model-the poly-alanine monomer model obtained from *Pf*TIM WT structure 2VFH was used for crystal 1 (asymmetric unit compatible with a monomer, 67% water content) while, the crystal 1 monomer was used for crystal 2 (asymmetric unit compatible with a dimer, 61% water content). To avoid model bias, loop-6 residues, water molecules, and ligand were deleted from the MR model and were added after few rounds of refinement on the basis of 2*F*_o_-*F*_c_ and *F_o_-F_c_* difference density maps contoured at 1σ and 3σ, respectively. Refinement of the structures was carried out using REFMAC (version 5.5.0109) [43]. Model building was done using COOT v0.7.2.1 [44] and superposition of structures was achieved by secondary superposition matching method in COOT [45]. The data collection and processing statistics and parameters after final round of model building and refinement are provided (Tables 2 and 3). While crystal-1 had an empty active site representing the *apo* form of the enzyme with loop-6 in its *open* conformation, crystal-2 (after DHAP soaking) showed an unmodeled blob at the active site and loop-6 in the *closed* conformation, Table S3. With careful examination of the crystal-2 unmodeled blob at the active site, a DHAP molecule was modeled with 0.6 and 0.8 fractional occupancy in subunits A and B, respectively. After achieving acceptable range of *R*_work_ and *R_free_* with model building and refinement (Table 3), the overall RMSD for structure superposition (SSM superpose) of *Li*TIM *apo* form monomer with *Pf*TIM WT monomer was ~1.7 Å and the RMSD for superimposition of the two subunits in the asymmetric unit of the *holo* form was ~0.5 Å. It was noted that overall RMSD for superposition of the *holo* forms of *Li*TIM dimer with that of the yeast TIM dimer (PDB ID: 1NEY) was high (~2 Å), likely due to the difference in the inter-subunit angle of their respective native dimers. All images showing structural details were prepared using Pymol version 1.2r1 [46]. Accessible and buried surface area and ΔG_dimer dissociation_ were determined using the PISA server [47] and using this ΔG, dissociation constant was calculated. Temperature (*B*) factors were analyzed by CCP4i version 2.2.1.

## ACKNOWLEDGMENT

Authors thank Dr. P. Vijayachari, Director, RMRC, ICMR, Port Blair, Andaman and Nicobar Islands, India, for providing the *Li* strain. VD and PRK acknowledge Vimal Raj Ratchagadasse and Vinod Kumar Kirubakaran’s (RMRC, ICMR, Port Blair, Andaman and Nicobar Islands, India) advice regarding *Li* culture maintenance. VP thanks Prof. Rikkert Wierenga (Biocenter Oulu, University of Oulu, Finland) for the kind gift of *tbtim* gene containing plasmid and Dr. Anne-Marie Lambeir for correspondence regarding *Tb*TIM purification and activity measurements. VP thanks Debarati Bandyopadhyay and Vivian J Roberts for assistance during *Tb*TIM expression, purification and initial activity measurements, Dr. Mamta Bangera for *Li*TIM crystal-2 data collection at the ESRF beamline, Sunitha Prakash for acquiring mass spectra, and James Paul and P F Babu at the MBU X-ray facility for their help. This work was part of VP and VD’s PhD thesis and they thank Prof Padmanabhan Balaram and Prof Mathur RN Murthy, Molecular Biophysics Unit, Indian Institute of Science, Bangalore, for their guidance and support for the work and helping with an help with an initial version of this MS.

## AUTHOR CONTRIBUTIONS

VP, VD, PRK and HB conceived the problem and designed the approach. VP and VD cloned and purified *Li*TIM. VD performed chaotrope induced denaturation and set up crystallization. VP carried out kinetic studies, temperature dependence of activity and stability, collected X-ray diffraction data and solved the structures with inputs from Mathur RN Murthy, MBU, IISc. VP VD and HB analysed and interpreted the data. VP and VD wrote the manuscript with inputs from all authors.

## Abbreviations

WHO: World Health Organization
TIM: Triosephosphate isomerase
*Li*: *Leptospira interrogans*
*Pf*: *Plasmodium falciparum*
*Tb*: *Trypanosoma brucei*
DHAP: Dihydroxyacetone phosphate
D-GAP: D-glyceraledhyde-3-phosphate
MG: Methylglyoxal
CD: Circular dichroism
SDS-PAGE: Sodium dodecyl sulfate polyacrylamide gel electrophoresis
LC-ESI/MS: Liquid chromatography coupled electrospray ionisation mass spectrometry
GdmCl: Guanidine hydrochloride

