## Supplementary figures and tables for "*Leptospira interrogans* triosephosphate isomerase: exploring the structural determinants of stability, high reaction rate and specificity" (from another version of this preprint: version 1, 2020-12-06)

**Supplementary Material**

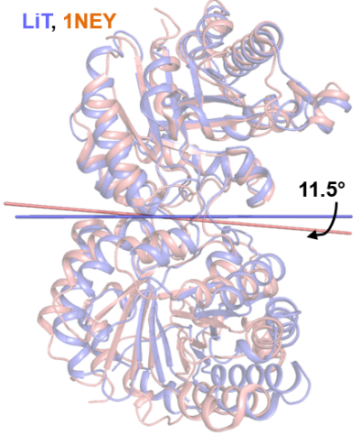

**Figure S1.** Overlay of LiTIM (PDB ID: 4YMZ) and Yeast TIM (PDB ID: 1NEY) dimer shows that the two subunits forming the native dimer are slightly differently oriented. Superposition was performed with SSM operation in Coot software using one subunit and allowing free movement of the other. The angle between the dimer axis found to be ~11.5°.

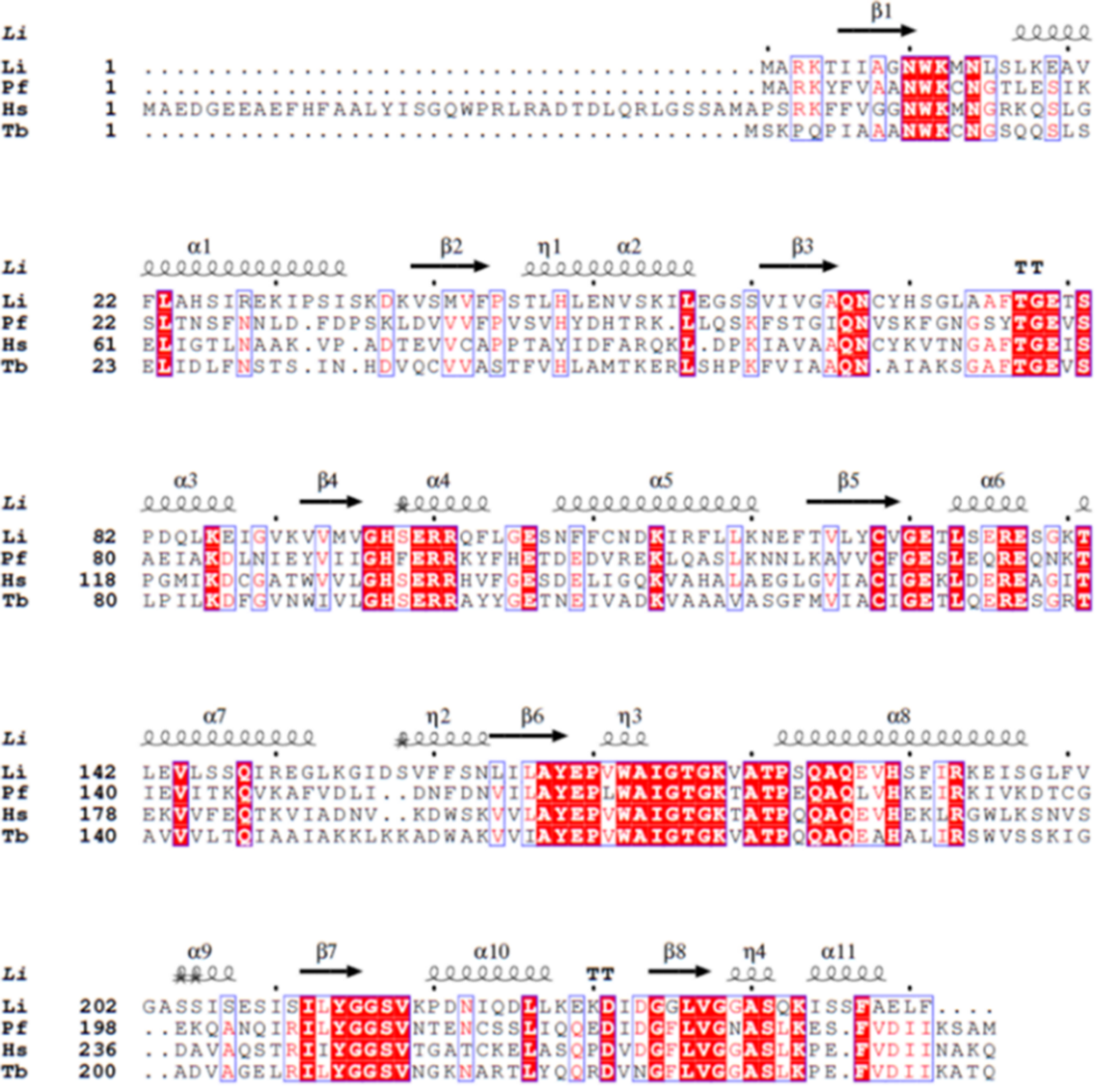

**Figure S2.** **Multiple sequence alignment of Human (Hs) TIM with TIM from the three pathogenic.** The TIM sequences of pathogenic species *Leptospira interrogans* (Li), *Plasmodium falciparum* (Pf), and *Trypanosoma brucei* (Tb) TIMs aligned with Human TIM sequence. The secondary structural elements denoted above the alignment were derived from the LiTIM structure. Alignment was performed with Clustal Omega software and the image was prepared using ESPript version3 (ESPript - http://espript.ibcp.fr).

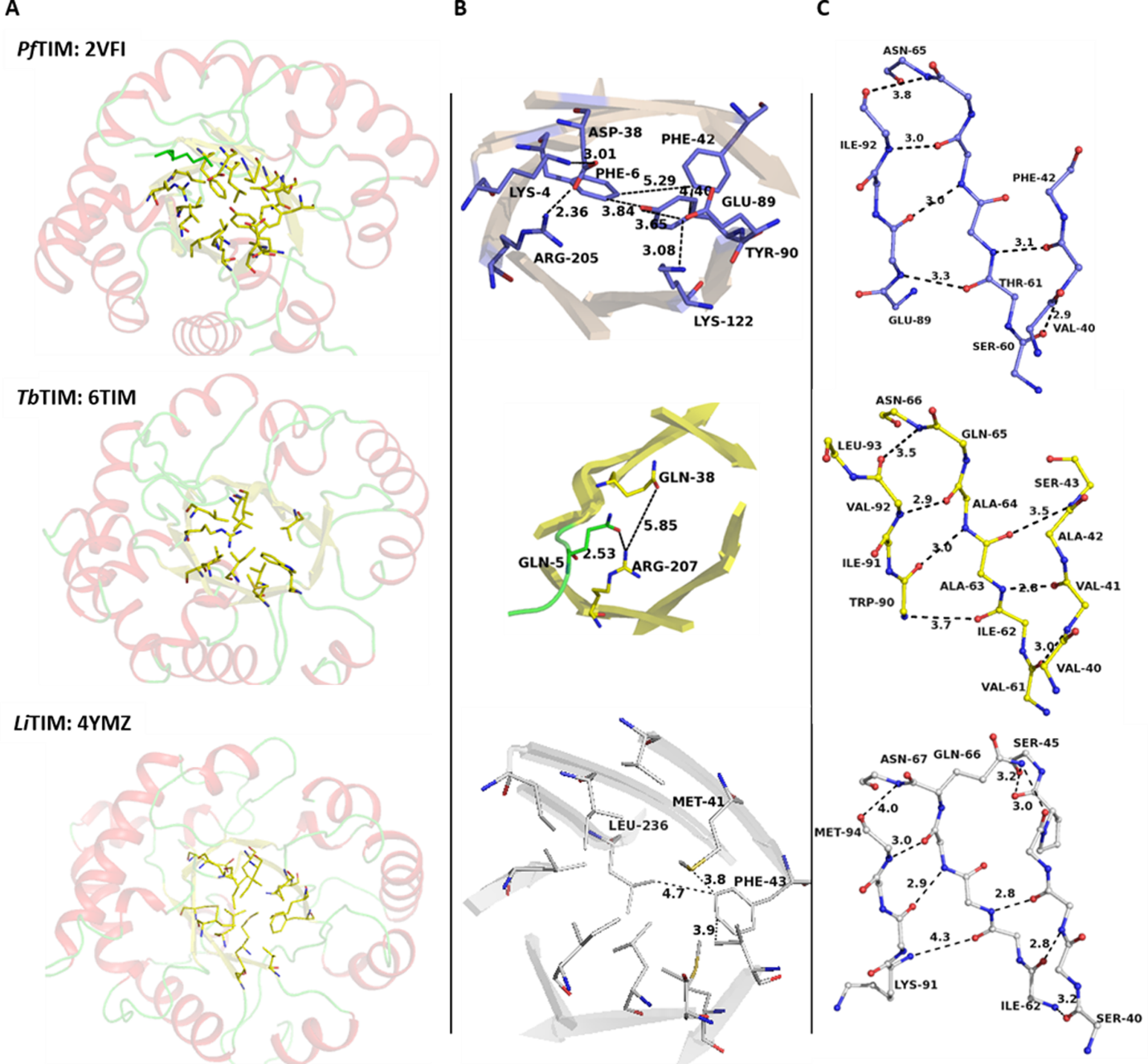

**Figure S3. The structural features of inner β- barrel core of LiTIM, PfTIM and TbTIM.** (**A**) Cartoon representation of TIM from *Plasmodium falciparum* (Pf), *Trypanosoma brucei* (Tb) and *Leptospira interrogans* (Li) with residue side chains pointing in the interior of the β- barrel core shown in sticks. (**B**) The β- barrel of PfTIM, TbTIM and LiTIM with residue side chains that show specific hydrophobic, aromatic or salt bridge interaction. More hydrophobic and charged residues line the barrel interior in PFTIM compared to TbTIm and LiTIM. (**C**) Backbone hydrogen bond interactions of β- strand 4 (residues 89- 92, PfTIM; 90- 93, TbTIM; and 91- 94, LiTIM) with β- strand 5. In PfTIM the strand registry stretches from residues 89- 92, with 4 hydrogen bonds, while, in LiTIM the two terminal hydrogen bonds are missing.

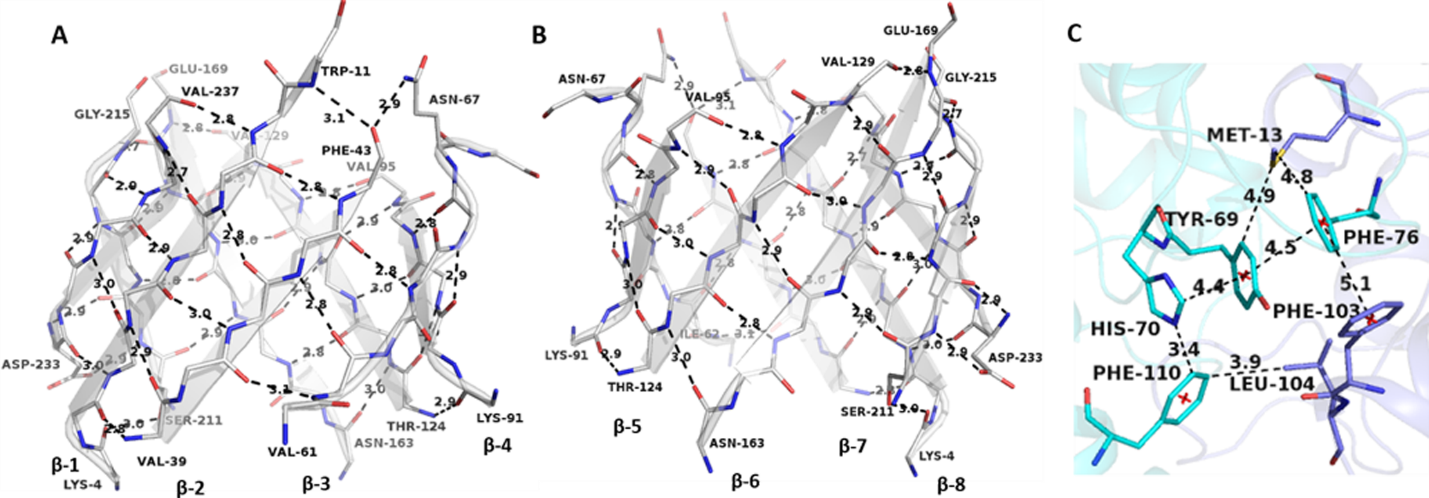

**Figure S4. The β-strand registry and dimer-interface aromatic cluster in LiTIM.** (**A**) and (**B**) showing the β-barrel core comprised of 8 parallel sheets in LiTIM structure. For clarity, in (A) and (B) only the backbone is shown in cartoon and stick representation. (**C**) Aromatic cluster at the dimer interface. The adjacent subunits are represented in blue and cyan colours.

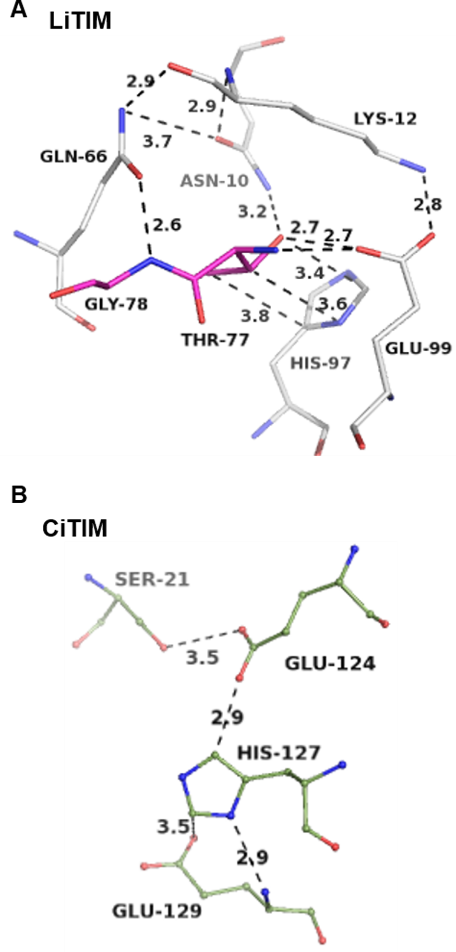

**Figure S5. Important interactions of catalytic residues K12, H97 and E99 in LiTIM and CiTIM.** (**A**) Dimer interface interactions involving T77 and G78 maintain the positioning of the active site residues K12, H97 and E99. Additionally, interactions involving residues Q66 and N10 provide additional stability. Residues of first subunit are shown in grey and that of the second subunit are shown in magenta. (**B**) Hydrogen bond network involving the catalytic residues H127 (H97 in LiTIM) in *Coccidioides immitis* (Ci) TIM (PDB ID: 3S6D). Presence of Glutamic acid on both sides of the imidazole ring may change the pKa of H127, also, may stabilize the ε^2^-H rotamer.

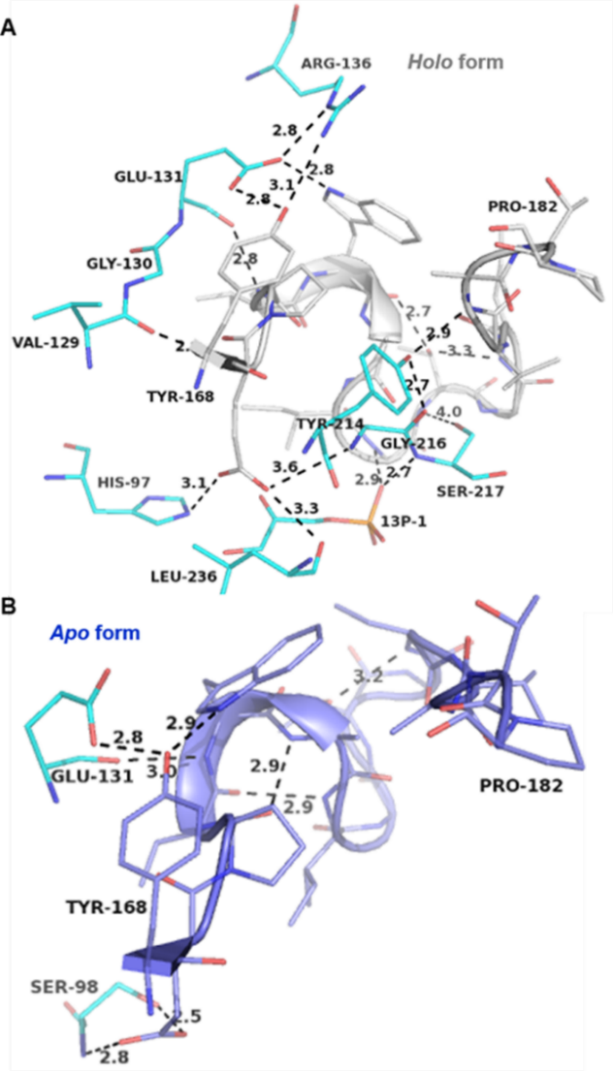

**Figure S6. Backbone and side chain hydrogen bonds involving Loop-6 residues in the *apo* and the *holo* form of LiTIM.** (**A**) Hydrogen-bonds made by loop-6 residues (grey) in the *holo* state with main body of the protein (cyan). These interactions contribute to stabilization of loop-6 *closed* conformation. (**B**) Hydrogen-bonds made by loop-6 residues (blue) in the *apo* state with main body of the protein (cyan). The lack of stabilizing network of H-bonds increases the conformational flexibility of the loop-6.

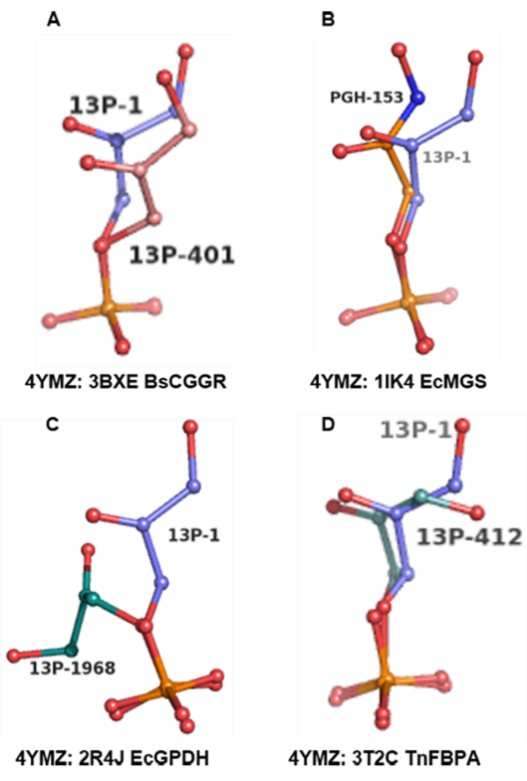

**Figure S7. Dihydroxyacetone phosphate (DHAP) binding modes in different enzyme active sites.** Superposition of DHAP bound in the active site of triosephosphate isomerase (blue, PDB ID: 4YMZ with (**A**) *Bacillus subtilis* cytoplasmic glycolysis gene regulator (salmon pink, PDB ID 3BXE); (**B**) *E. coli* methylglyoxal synthase (PDB ID 1IK4); (**C**) *E. coli* glycerol-3-phosphate dehydrogenase (deepteal, PDB ID 2RJ4) and (**D**) *Thermoproteus neutrophilus* fructose-1,6- bisphosphate aldolase/phosphatase (green cyan, PDB ID 3T2C). Superposition was perfomed by the pairfitting function in Pymol 1.2r1 using the phosphate atoms of DHAP.

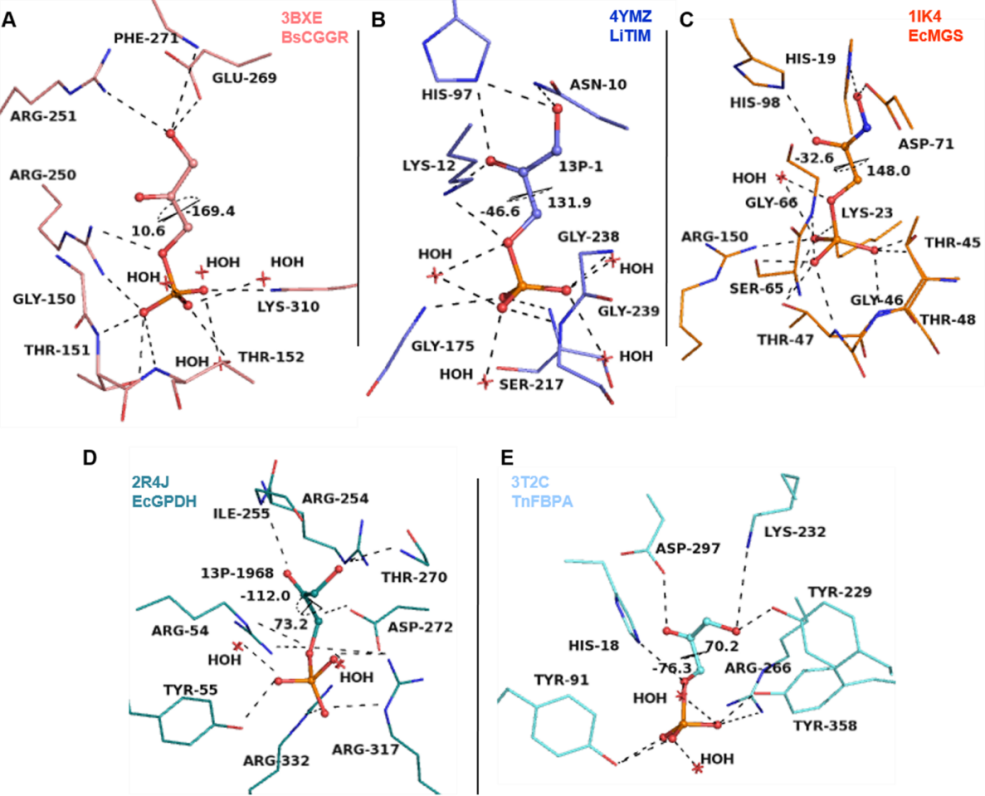

**Figure S8. DHAP Binding mode and interactions with the residues lining the active site pocket of different enzymes.** DHAP conformation, binding strength is determined by the active site residues and their geometry which is specific to each enzyme. (**A**) *Bacillus subtilis* central glycolytic gene regulator, and (**B**) *Leptospira interrogans* TIM active site, (**C**) The transition state analogue phosphoglycolohydroxamate (PGH) in *E. coli* methylglyoxal synthase, MGS, active site, (**D**) DHAP in the active site of *E. coli* α-glycerophosphate dehydrogenase and (**E**) The respective PDB IDs are indicated on the figure. Plausible hydrogen bonds are indicated as dashes, ligand is shown as ball and stick, water molecules are shown as red crosses and active site residues are shown as sticks. The dihedral angle values for C1C2C3O3 and O2C2C3O3 are indicated on the figures. All figures were prepared using Pymol 1.2r1.

**Table S1. Primers used for LiTIM cloning.**

| Primer name | Primer sequence (5'-3') | Restriction site |
| --- | --- | --- |
| LiTIMFP | CATGCCATGGCTCGTAAGACAATTATTGCAGGAAACTGG (Tm-81.3˚C) | NcoI |
| LiTIMRP | GCGGGATCCTTAGAAAAGTCCAGCGAAGGAACTG (Tm-79.6˚C) | BamHI |

**Table S2. Crystallization conditions under which the LiTIM crystals were obtained.**

|  | Crystal 1 | Crystal 2 |
| --- | --- | --- |
| Growth time | ~35 days | ~25 days |
| Buffer | HEPES (100mM) pH7.5 | BIS-TRIS (100mM) pH6.6 |
| Precipitant | PEG 6K 18% | PEG 8K 16% |
| EDTA | 1mM) | 10mM) |
| Salt | CaCl₂ (20mM) | (NH_4_)₂SO₄ (1mM) |
| Sodium azide | 1mM | 1mM |
| Protein: mother liquour | 3μl (9mg/ml) :3μl | 2.5μl (15mg/ml) :2.5μl |
| Crystal soaking | No | In 280mM DHAP |
| Cryoprotectant | 10% Ethylene glycol | 15% Ethylene glycol |

**Table S3. Data collection and processing statistics for LiTIM crystal1 (*apo* form) and Crystal2 (*holo* form).**

|  | Crystal 1 | Crystal 2 |
| --- | --- | --- |
| Space group | *P*3_1_21 | *P*3_2_21 |
| PDB ID | 4X22 | 4YMZ |
| Cell parameters  a (Å), b (Å), c (Å)  α (°), β (°), γ (°) | 103.0, 103.0, 66.1  90.0, 90.0, 120.0 | 103.8, 103.8, 113.5  90.0, 90.0, 120.0 |
| Resolution range (Å) | 44.60 - 2.08 | 56.75 - 1.87 |
| Average mosaicity (˚) | 0.77 | 1.10 |
| Overall *R*_merge_(%)^#^ | 0.055 | 0.118 |
| Overall Completion (%) | 95.9 | 92.7 |
| Overall <I>/<σI> | 25.9 | 13.3 |
| # of observed reflections | 208090 | 647205 |
| # of unique reflections | 23354 | 54809 |
| Overall Multiplicity | 8.9 | 11.8 |
| Wilsons B factor (Å^2^) | 22.4 | 22.8 |
| Starting MR model | 2VFH | 4X22 |
| # of subunits/ asymmetric unit | 1 | 2 |

^#^*R*_merge_= Σ*_hkl_*Σ*_i_│I_i_*(*hkl*) *–* (*I*(*hkl*))*│*/Σ*_hkl_*Σ*_i_I_i_*(*hkl*), where *I_i_*(*hkl*)is the *i*_th_ observation of *I*(*hkl*) and *I*(*hkl*) is its mean intensity.

**Table S4. Data refinement statistics for LiTIM crystals.**

|  | Crystal 1 | Crystal 2 |
| --- | --- | --- |
| Number of used reflections | 22141 | 52007 |
| % observed | 95.1 | 92.8 |
| *R*_work_(%) | 15.4 | 17.1 |
| *R*_free_(%) | 19.1 | 21.1 |
| FOM | 89.7 | 88.2 |
| Model quality:  Number of protein atoms | 1874 | 3739 |
| Number of water molecules | 221 | 308 |
| Number of ligand atoms | 24 | 90 |
| Matthews coefficient | 3.71 | 3.23 |
| Average B-factor (Ǻ^2^): | 23.0 | 27.8 |
| RMS deviation from ideal:  Bond length (Ǻ)  Bond angle (°)  Chiral volume (Ǻ^3^) | 0.02  1.89  0.12 | 0.02  2.01  0.13 |
| Ramachandran statistics:  Favored region (%)  Allowed region (%)  Disallowed region (%) | 97  3  0 | 97  3  0 |

**Table S5. The backbone dihedral angles (φ and ψ) and the normalized temperature factors (*B*) for the loop-6 residues in the *apo* and the *holo* form of LiTIM.**

| Residue position | *Apo* form | | | *Holo* form | | |
| --- | --- | --- | --- | --- | --- | --- |
|  | **φ (°)** | **ψ (°)** | ***B*_nor_^#^** | **φ (°)** | **ψ (°)** | ***B*_nor_^#^** |
| **Y168** | - | 120 | 0.8 | - | 115 | 0.8 |
| **E169** | −103 | 115 | 0.8 | −133 | 95 | 1.0 |
| **P170** | −82 | 75 | 0.9 | −64 | 34 | 1.0 |
| **V171** | −172 | −33 | 1.0 | −63 | −35 | 1.1 |
| **W172** | −54 | −29 | 1.1 | −58 | −14 | 1.2 |
| **A173** | −102 | 1 | 1.8 | −105 | −4 | 1.1 |
| **I174** | −91 | 99 | 2.4 | −87 | 109 | 1.0 |
| **G175** | 51 | 44 | 3.1 | 76 | 0 | 1.2 |
| **T176** | −112 | 3 | 2.7 | −92 | −12 | 1.4 |
| **G177** | −97 | 69 | 2.8 | 80 | 0 | 1.7 |
| **K178** | −62 | 154 | 2.7 | −105 | 82 | 1.5 |
| **V179** | −131 | 130 | 2.5 | −66 | 140 | 1.2 |
| **A180** | −96 | 130 | 1.8 | −59 | 141 | 1.1 |
| **T181** | −85 | 163 | 1.4 | −67 | 159 | 1.2 |
| **P182** | −61 | −31 | 1.3 | −51 | −45 | 1.2 |

^#^ For normalization, the average main chain temperature factor for a residue was divided by the overall average of main chain temperature factor for the whole chain.

**Table S6. Dihedral angles of bound substrate in the five reported class of enzymes. Atom numbering is as per given in Figure 8.**

| DIHEDRAL ANGLE (°) | 4YMZ  LiTIM | 1IK4  EcMGS | 3BXE  BsCGGR | 2R4J  EcGPDH | 3T2C  TnFBPA |
| --- | --- | --- | --- | --- | --- |
| **O1C1C2C3** | 171.2 | 179.3 | 167.8 | 101.9 | 30.7 |
| **C1C2C3O** | 131.9 | 148.0 | −169.4 | 73.2 | 70.2 |
| **C2C3O3P** | −178.0 | 177.0 | −154.3 | −62.6 | 167.1 |
| **O1C1C2O2** | −10.5 | 0.0 | −12.2 | −73.0 | 177.3 |
| **O2C2C3O** | −46.6 | −32.6 | 10.6 | −112.0 | −76.3 |

**Table S7. The distance of different atoms of a DHAP molecule after overlaying the molecules from different structures as mentioned in the table. The phosphate atoms were used for the overlay and the rest of the molecule was allowed to move.**

| DISTANCE (Å) | 4YMZ:1IK4 | 4YMZ:3BXE | 4YMZ:2R4J | 4YMZ:3T2C |
| --- | --- | --- | --- | --- |
| **O1:O1** | 1.7 | 2.9 | 7.8 | 2.1 |
| **C1:C1** | 1.1 | 2.4 | 5.7 | 0.9 |
| **C2:C2** | 0.5 | 1.9 | 3.1 | 0.8 |
| **O2:O2** | 0.7 | 1.8 | 3.3 | 0.8 |
| **C3:C3** | 0.2 | 1.6 | 1.7 | 0.7 |

**Table S8. The phosphate oxygen mediated potential hydrogen bonds made by substrate (DHAP) in different active site pockets.**

| PDB ID | Neutral H-bonds | Charged H-bonds |
| --- | --- | --- |
| **4YMZ** | Backbone NH-4; SerOH- 1 | Lys-1 |
| **1IK4** | Backbone NH-4; ThrOH-3; SerOH-1 | Arg-1, Lys-1 |
| **3BXE** | TyrOH-1; ThrOH-1 | Arg-3 each with 3 H-bonds |
| **2R4J** | TyrOH-3; | His-1, Arg-1 |
| **3T2C** | Backbone NH-2; ThrOH-2 | Arg-2, Lys-1 |

**Table S9. Natural variation observed at positions, 10, 14 and 94 (N/N/M triad combination for LiTIM) in the curated dataset of TIM sequences (residue numbering as in LiTIM sequence).**

| Organism  (#)*^a^* | Residue position (η)*^b^* | Residue variability and respective occurrence frequency*^c^* | | | | |
| --- | --- | --- | --- | --- | --- | --- |
| Bacteria  (5216) | 10 (3) | N 5029 | S 184 | G 3 |  |  |
|  | 14 (15) | N 3805 | H 815 | Y 234 | A 201 | F 140 |
|  | 94 (9) | I 2861 | L 1211 | V 762 | E 212 | A 77 |
| Eukaryote  (819) | 10 (3) | N 813 | S 5 | V 1 |  |  |
|  | 14 (9) | N 799 | - 5 | S 3 | H 3 | C 2 |
|  | 94 (5) | I 638 | L 109 | V 62 | M 5 | E 5 |
| Archaea  (241) | 10 (1) | N 241 |  |  |  |  |
|  | 14 (2) | N 799 | - 5 | S 3 | H 3 | C 2 |
|  | 94 (3) | L 213 | I 23 | M 6 |  |  |

*^a^* total number of sequences used in the analysis in the respective subgroup of organisms. *^b^*η is the total number of different amino acids observed at a position. *^c^* 5 most frequently occurring amino acids with respective frequencies observed in decreasing order.
